# Catalysis of cataract-associated human γD crystallin aggregation via dynamic disulfide exchange

**DOI:** 10.1101/300608

**Authors:** Eugene Serebryany, Shuhuai Yu, Sunia A. Trauger, Bogdan Budnik, Eugene I. Shakhnovich

**Affiliations:** Department of Chemistry and Chemical Biology, Harvard University, Cambridge, MA; State Key Laboratory of Food Science and Technology, Jiangnan University, Wuxi, Jiangsu, China; Harvard FAS Science Core Facility, Cambridge, MA

## Abstract

Several mutations in human γD-crystallin (HγD), a long-lived eye lens protein, cause misfolding and aggregation, leading to cataract. Surprisingly, wild-type HγD catalyzes aggregation of its cataract related W42Q variant while itself remaining soluble – the inverse of the classical prion-like scenario whereby misfolded polypeptides catalyze aggregation of natively folded ones. The search for a biochemical mechanism of catalysis of W42Q aggregation by WT has revealed that WT HγD can transfer a disulfide bond to the W42Q variant. The transferred disulfide kinetically traps an aggregation-prone intermediate made accessible by the W42Q mutation, facilitating light-scattering aggregation of the W42Q variant. The aggregating variant thus becomes a disulfide sink, removing the disulfides from solution. Such redox “hot potato” competitions among wild-type and mutant or modified polypeptides may be relevant for many long-lived proteins that function in oxidizing environments. In these cases aggregation may be forestalled by inhibiting disulfide flow toward damaged polypeptides.

## Introduction

*In vivo* populations of many proteins exhibit conformational and chemical heterogeneity (Bah and Forman-Kay, 2016; Deis et al., 2014; Fuchs et al., 2011; Hornbeck et al., 2012; Lu et al., 1998; Murzin, 2008; Walsh et al., 2005; Xin and Radivojac, 2012). It may arise from post-translational modifications; somatic mutations; roughness of the conformational landscape; or simply as a natural effect of heterozygosity. Biochemical heterogeneity is particularly important for highly abundant, long-lived (low-turnover) proteins such as the collagens and aggrecans of the joints; elastins of the skin; and crystallins of the eye, which accumulate damage throughout the course of life (de Graff et al., 2016; Ma et al., 1998; Truscott et al., 2016). For well-structured proteins, partially populated intermediate and misfolded conformational states give rise to heterogeneous conformational ensembles in living cells, which sometimes leads to aggregation (Bartlett and Radford, 2009; Chiti and Dobson, 2009; Horwich, 2002; Wang et al., 2016). The emergent properties of subtly heterogeneous macromolecular populations are largely unexplored, yet there is ample evidence that they can be significant (Bah and Forman-Kay, 2016; Forman-Kay and Mittag, 2013; Walsh et al., 2005). Perhaps the most spectacular case is the prion effect, wherein polypeptides in an aggregation-prone conformational state can catalyze conformational conversion and aggregation in the natively-folded polypeptides (Aguzzi and Calella, 2009). Specific mutations or post-translational modifications can enhance or suppress these aggregation-promoting interactions, e.g., in the ALS-associated enzyme superoxide dismutase 1 (Grad and Cashman, 2014); in β_2_-microglobulin, associated with dialysis amyloidosis (Karamanos et al., 2014); and the yeast prion Sup35 (Tessier and Lindquist, 2007).

Intramolecular disulfides are crucial for the correct folding of many proteins that function in relatively oxidizing environments, such as antibodies and hormones in the blood, transmembrane proteins, digestive enzymes, as well as a wide variety of protein toxins (Creighton, 1988; Fass and Thorpe, 2018; Go et al., 2015; Sevier and Kaiser, 2002; Thornton, 1981; Trivedi et al., 2009; Woycechowsky and Raines, 2000). At the same time, internal disulfides are well-known conformational traps on the protein folding landscape, capable of stabilizing both productive folding intermediates and aggregation-prone misfolded states (Arolas et al., 2006; Chang et al., 2001; Hua et al., 2002; Serebryany et al., 2016b; Toichi et al., 2013). Redox chemistry is known to be crucial for many proteins’ aggregation pathways. These include superoxide dismutase 1 (SOD1), associated with amyotrophic lateral sclerosis (Toichi et al., 2013), as well as β_2_-microglobulin, the amyloid-forming protein in dialysis amyloidosis (Chen and Dokholyan, 2005; Smith and Radford, 2001). Disulfide scrambling is an important failure mode of therapeutic antibodies (Trivedi et al., 2009) and may be involved in light-chain amyloidosis (Connors et al., 2007). Domain-swapping is a particularly sensible mechanism for aggregation via disulfide-stabilized intermediate states, and indeed this may play a role in serpinopathies (Ronzoni et al., 2016). Dynamic disulfide exchange among identical or nearly identical polypeptides has been only recently recognized, in the case of protein disulfide isomerase (Oka et al., 2015), and is likely to be found in other systems. Most known proteins are capable of undergoing some reversible post-translational modifications, including a wide variety of disulfide bonds (Bah and Forman-Kay, 2016; Cook and Hogg, 2013; Walsh et al., 2005). Redox potential in various tissues changes over time, in the course of development or across the cell cycle, due to episodes of oxidative stress, or in the course of aging (Banerjee, 2012; Jones, 2010; Sarsour et al., 2009).

The eye lens is a remarkable study in proteome aging, since the proteins in its core are never replaced. Their eventual misfolding and aggregation leads to cataract disease – lens opacification due to light scattering by the protein aggregates (Moreau and King, 2012b). Crystallins in the lens core are highly stable and soluble, resisting any long-range packing interactions, including formation of both native (crystals) and non-native (amyloid or amorphous) aggregates (Serebryany and King, 2014). However, they accumulate damage over time, such as Cys and Trp oxidation, as well as deamidation, truncation, and other changes (Hains and Truscott, 2007, 2008; Hoehenwarter et al., 2006; Lampi et al., 2014). One of the most readily formed yet most consequential post-translational modifications that accumulate with aging is formation of disulfide bonds. Cells of the human eye lens core lack nuclei or organelles and are metabolically quiescent (Bloemendal et al., 2004; Wride, 2011), so their cytoplasm gets more oxidizing over time (Friedburg and Manthey, 1973). Due to this lack of active metabolism, neither the redox enzymes, nor the crystallins, which make up the bulk of the lens proteome, turn over (Michael and Bron, 2011; Shang and Taylor, 2004). The result is progressively higher disulfide content in lens crystallins during aging (Giblin et al., 1995; Michael and Bron, 2011; Ozaki et al., 1987; Spector and Roy, 1978; Yu et al., 1985). Oxidation of specific Cys residues to disulfides in lens crystallins correlates with the onset and progression of cataract disease (Cherian-Shaw et al., 1999; Fan et al., 2015; Takemoto, 1997a, b).

We recently discovered a surprising interaction between the highly stable wild-type human γD-crystallin (HγD) and its partially destabilized, oxidation-mimicking W42Q mutant, whereby the WT protein promoted aggregation of the mutant without itself aggregating (Serebryany and King, 2015). The W42Q mutation mimics oxidative damage to Trp side chains (conversion to kynurenine, which is more hydrophilic) that is known to arise during aging (Hains and Truscott, 2007; Hoehenwarter et al., 2006; Moran et al., 2013); the W42R variant, whose biophysical properties are highly similar, causes hereditary cataract in humans (Ji et al., 2013; Serebryany et al., 2016a; Serebryany et al., 2016b; Wang et al., 2011).

The “inverse-prion” aggregation phenomenon may represent a new class of protein-protein interactions emerging in aging, damaged, heterozygous, or otherwise heterogeneous proteomes. Here we addressed the mystery of the “inverse-prion” catalysis of mutant aggregation by WT and discovered that the WT protein can function as an oxidoreductase: By oxidizing the mutant proteins, WT HγD triggers a chain of events leading to aggregation of the mutant variants. Specifically, a pair of Cys residues in the C-terminal domain of HγD is capable of forming a labile disulfide bond. These disulfides are dynamically exchanged among crystallin molecules under physiological conditions in solution, creating a “redox hot potato” competition: the disulfide is passed around until it is transferred to a destabilized variant where it can lock the aggregation-prone intermediate; then aggregation ensues, taking both the destabilized variant and the disulfide out of solution.

## Results

### WT HγD crystallin must be oxidized in order to catalyze aggregation of the W42Q mutant

We have previously reported (Serebryany and King, 2015) the surprising interaction between wild-type HγD crystallin and its cataract-associated W42Q mutant, whereby the WT protein promoted the mutant’s aggregation in a temperature- and concentration-dependent manner without aggregating itself. We term this catalytic-like effect of the WT protein as “inverse-prion” activity. In a separate study we found that formation of an internal disulfide bond was crucial for the mutant protein’s ability to aggregate (Serebryany et al., 2016b). Therefore, we explored the possibility that redox chemistry is involved in the observed inverse-prion aggregation-promoting mechanism.

We found that fully reduced purified WT protein was inactive as an inverse prion. However, it could be converted to the active form *in vitro* (**Figure 1B)** by a commonly used oxidizing treatment with a 1:3 mixture of copper(II) and phenanthroline (Careaga and Falke, 1992; Kobashi, 1968). Reduction of the WT_ox_. sample with tris(2-carboxyethyl)phosphine (TCEP) inactivated it again (**Figure 1B**). Isotopically resolved whole-protein electrospray mass spectrometry confirmed that the peak isotope mass of the WT sample prior to oxidation was 20,605.9 Da., which precisely matches the weight predicted from the amino acid sequence (20,605.8 Da.), and the overlap of isotopic distributions was also nearly perfect between the theoretical and experimentally determined mass spectra (**Figure 1C**). The same experiment was carried out on the oxidized sample whose activity was assayed in 1B, and the resulting molecular weight and isotope distribution both precisely matched those predicted for the protein in the case of one intramolecular disulfide bond per polypeptide ( –SH + –SH –> –S–S–, hence the loss of 2 Da.) following the oxidizing treatment. Thus, the oxidation was highly specific, leaving the remaining four Cys residues unaffected, and essentially quantitative under these conditions.

**Figure 1:**
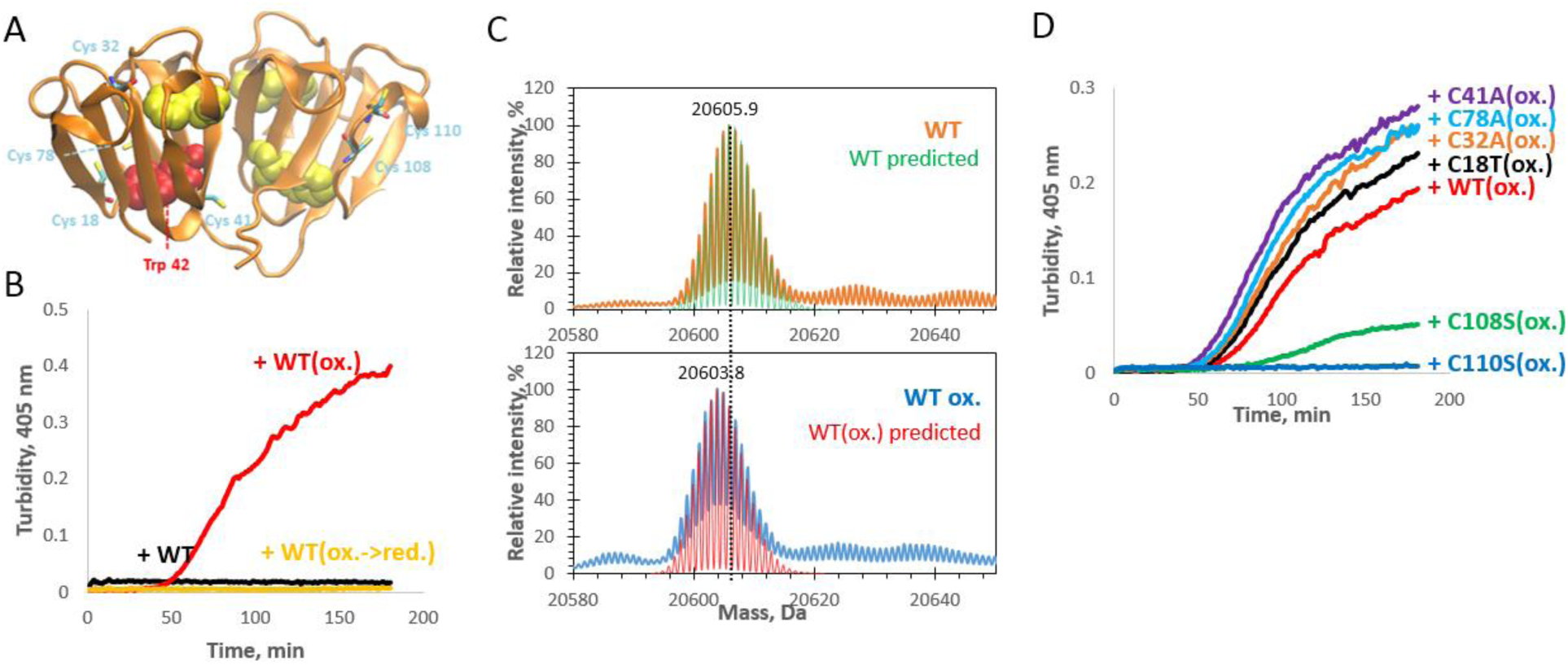
WT HγD can be toggled between the inert and inverse-prion states *in vitro* by formation or reduction of the 108-SS-110 internal disulfide bond. (A) Representation of the WT crystal structure in orthosteric projection (PDB ID 1HK0) showing the location of the six Cys residues, as well as Trp42. (B) Turbidity traces from mixtures of reduced W42Q with reduced (black), oxidized (red), and oxidized and re-reduced WT (yellow). Concentration of each protein was 50 μM. (C) Whole-protein isotopically resolved mass spectra showing WT(ox.) to be 2 Da. lighter than the reduced version, indicating formation of one internal disulfide bond per molecule. Theoretical isotopically resolved mass spectra based solely on the atomic composition (*green*, *red*) are overlaid on the respective experimentally determined deconvoluted isotopically resolved mass spectra (*orange*, *blue*). (D) WT HγD and variants at every Cys position were oxidized in parallel, and their ability to catalyze W42Q aggregation was tested (each protein at 50 μM). Resulting turbidity traces confirmed Cys108 and Cys110 as the residues predominantly responsible for inverse-prion activity.

### A specific disulfide bond in the WT protein generates inverse-prion activity

The native-state crystal structure of HγD (**Figure 1A**) has no disulfide bonds, despite the statistically high proportion of Cys residues in this protein (6 out of 173 residues). Tissue proteomic studies have revealed disulfide formation in this protein in the oxidizing cytosol of aged and cataractous lenses (Fan et al., 2015; Hanson et al., 1998). At least two pairs of Cys residues (18-78 and 108-110) are sufficiently close in the native structure that a disulfide bond might form with only a short-range structural perturbation. Other disulfides have been observed following mutation or denaturation of the protein (Hanson et al., 1998; Serebryany et al., 2016b).

We therefore used site-directed mutagenesis to eliminate, one at a time, each of the six Cys residues on the WT background. Simultaneous oxidation of WT and Cys-mutant proteins under identical conditions showed no declines in inverse-prion activity, relative to WT, for any mutant except C108S and C110S, which lost all or most of this activity (**Figure 1D**). Thus, the 108-SS-110 disulfide appeared to be crucial for the inverse-prion activity. We were able to directly detect peptides containing this disulfide by LC/MS/MS of chymotryptic digests of the oxidized protein (**Table SI1**). This was additionally confirmed by top-down proteomics with incomplete chymotrypsin/GluC digestion, which yielded a peak at 10,382.4101 Da. (consistent with residues 65-150 with one disulfide), which fragmented to yield a peak at 6921.9584 Da. (consistent with residues 93-145 with 108-SS-110, as no other Cys residues are present between these positions). By contrast, combined tryptic/chymotryptic digests revealed that the N-terminal Cys residues (particularly Cys18 and Cys78) were mostly in the reduced state (**Table SI1**). A smaller amount of Cys18-Cys32 disulfides was observed, although detection of Cys32- and Cys41-containing peptides proved challenging, in line with previous proteomic data (Hains and Truscott, 2008).

Notably, isotopically resolved mass spectrometry of the C108S and C110S variants following oxidative treatment revealed significant fractions of each variant with a +16 Da. mass shift, suggesting a gain of one Oxygen atom (**Figure SI 2**). The most common site of such a modification in proteins is a Met residue, although stable adducts can also form at Trp, Tyr, or His side chains. Thus, in our experiments the 108-SS-110 bond in HγD appeared to act as a preferred oxidation scavenger site to avoid irreversible oxidation of other residues. Furthermore, the C108S/C110S double mutant exhibited much less of the +16 Da. species (**Figure SI 2**), suggesting that this form of covalent damage was cysteine-mediated in some way.

### Disulfide transfer from WT to W42Q promotes W42Q aggregation

We next tested whether this newly discovered oxidoreductase activity was related to the inverse-prion activity of the WT protein. This was accomplished by incubating a 1:1 mixture of oxidized WT and reduced W42Q until it turned turbid, indicating insolubilization of some fraction of the latter. Then, fresh W42Q was added and the reaction repeated for a total of four cycles, so that by the end the total amount of W42Q added was roughly 4x WT. By the fourth cycle, turbidity was visibly lower than initially. After each step, the turbid solution was centrifuged, and the pelleted fractions washed, resuspended in sample buffer, and combined.

Whole-protein isotopically resolved ESI-TOF (**Figure 3**) revealed that the supernatant at the end of the fourth aggregation cycle contained predominantly reduced WT protein and reduced W42Q. Thus, most of the disulfide-containing WT protein had been converted to the reduced form. Analysis of the pelleted fraction proved more difficult, since the proteins were in an aggregated state. The aggregates were partially resolubilized in low-pH buffer (pH 5 ammonium acetate). ESI-TOF of this resolubilized protein revealed that it consisted predominantly of W42Q with one disulfide bond (-2 Da. mass shift), consistent with our initial observations (Serebryany and King, 2015). It follows that oxidized WT HγD promoted W42Q aggregation by transferring a disulfide to it. The result can be summarized as a chemical reaction where the reactants WT(ox.) and W42Q are both soluble, while the products are soluble WT and insoluble W42Q(ox.). Aggregation of the oxidized mutant may shift the equilibrium in favor of disulfide transfer to the mutant.

**Figure 3:**
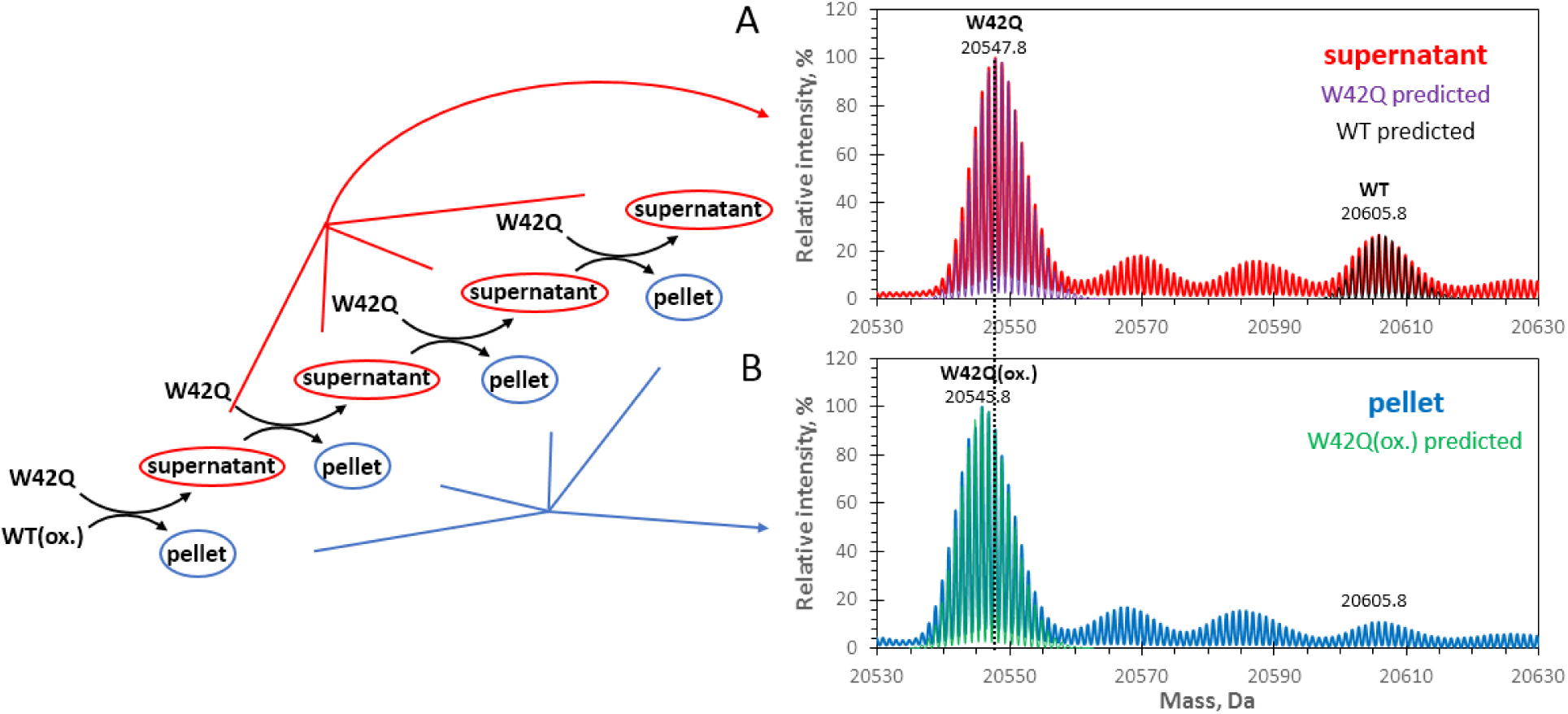
Disulfide transfer from soluble WT(ox.) to soluble W42Q generates soluble WT and aggregated W42Q(ox.) Whole-protein isotopically resolved mass spectra for the supernatant (A) and pelleted precipitate (B) fractions from aggregation mixtures of WT(ox) with W42Q are shown. As shown in the schematic (*left*), W42Q was added at each step, but WT(ox.) only initially, hence total final [W42Q] in the experiment was four times the initial [WT(ox.)]. The pooled supernatant fractions (A) clearly contained only reduced W42Q, as evidenced by the exact match between the observed (*red*, 20547.8 Da.) and theoretically predicted (*purple*) isotope distributions, and the WT(ox.) sample had been almost completely reduced back to WT, since the observed peak (*red*, 20,605.8 Da.) skewed only slightly to the left compared to the theoretically predicted isotope distribution for the fully reduced WT (*black*). The pooled pelleted fractions contained exclusively −2Da. shifted W42Q, as evidenced by the exact overlap between the experimental data (*blue*, 20545.8 Da.) and theoretically predicted isotope distribution for W42Q with one internal disulfide per molecule (*green*). Hence, the internal disulfide was transferred from soluble, oxidized WT to the soluble reduced W42Q during the course of the reaction, leading to formation of insoluble, oxidized W42Q.

The ability of W42Q to accept a disulfide bond and aggregate depended on the nature of the oxidant disulfide. Thus, glutathione disulfide (GSSG) caused strong aggregation of W42Q with comparable kinetics to those observed upon addition of the oxidized WT protein. By contrast, the same amount of hen egg lysozyme had no effect on W42Q (**Figure 4**), despite the fact that lysozyme contains four disulfide bonds per molecule. We interpret the difference as arising from the much higher accessibility and/or redox potential of both the reversible Cys108-Cys110 disulfide in WT HγD and the GSSG disulfide as compared to the structural disulfides in lysozyme. It is worth noting that disulfide geometry has a strong effect on whether it is a catalytic or a structural bond (Cook and Hogg, 2013), and that the close sequence-proximity of Cys108 and Cys110 is likely to give it a strained, catalytic-like configuration (Cook and Hogg, 2013; Fass and Thorpe, 2018; Woycechowsky and Raines, 2003).

**Figure 4:**
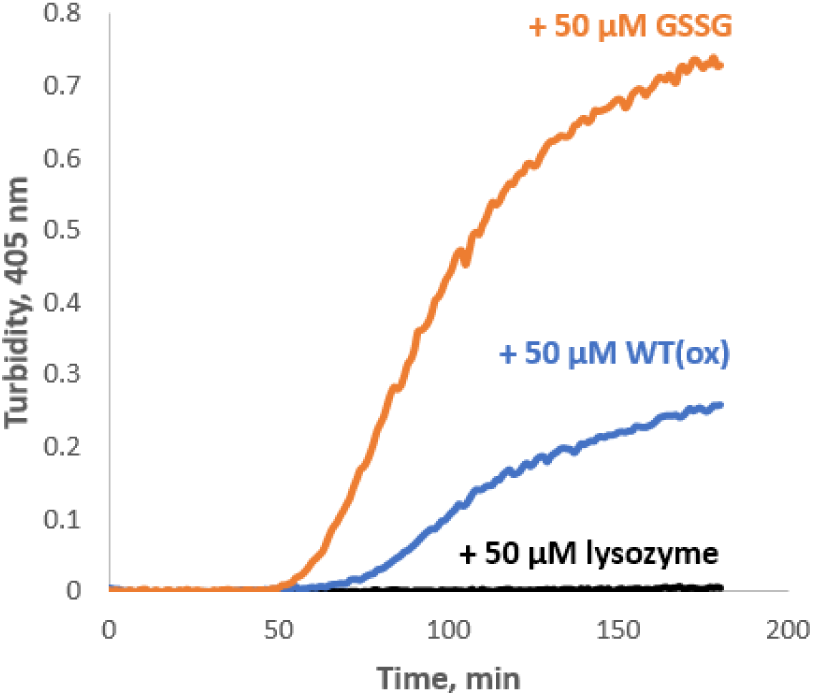
W42Q aggregation depends on dynamic nature of the oxidant disulfide. 50 μM W42Q protein was incubated at 37 °C and pH7 in the presence of 50 μM hen egg lysozyme (*black*), 50 μM WT(ox) (*blue*), or 50 μM oxidized glutathione (*orange*). The four disulfide bonds per mole in lysozyme are not dynamic, in contrast to the single disulfide per mole of oxidized WT or oxidized glutathione.

The disulfide required for W42Q aggregation was distinct from the dynamic 108-SS-110 disulfide in WT. LC/MS/MS (**Table SI 3**) revealed the internal disulfide in the W42Q aggregates to be predominantly 32-SS-41, consistent with previous results (Serebryany et al., 2016b) as well as tissue proteomic analysis (Fan et al., 2015). Accordingly, the W42Q/C110S double-mutant aggregated comparably to W42Q at neutral pH when mixed with the WT(ox) protein (**Figure SI 4**). The ability to form the 108-SS-110 disulfide, however, appeared to confer modest aggregation resistance at lower concentrations of the oxidizing agent glutathione disulfide (GSSG), perhaps because this site acted as an oxidation decoy, reducing or delaying oxidation of the N-terminal domain. This resistance, however, appeared to come at the cost of increased aggregation propensity at higher oxidant concentrations (**Figure SI 5**). It is possible that 108-SS-110 formation reduced solubility of the monomer or reversibility of the aggregated state (perhaps via partial intermolecular crosslinking). We also cannot rule out that 108-SS-110 in the mutant promotes formation of the N-terminal disulfide under some conditions. Indeed, lower pH (pH 5) led to a dramatically different behavior for the W42Q and W42Q/C110S mutants. When mixed with the oxidizing agent copper(II)/phenanthroline, only the single-mutant aggregated at pH 5, whereas both did so at pH 7 (**Figure SI 6**). This difference suggests that Cys110 may retain its reactivity even at reduced pH; however, the mechanism of aggregation, including any potential intramolecular disulfide transfer, under these conditions remains to be investigated.

Since resolubilization of the aggregated state under mild conditions was incomplete, we also resolubilized them in 5% SDS at 95 °C, pH5, and quantified the distribution of the number of free thiol groups per molecule using a PEGylation gel shift assay similar to those reported in (Basilio et al., 2014; Goodson and Katre, 1990; Lu and Deutsch, 2001). Free thiol groups reacting with maleimide-conjugated polyethylene glycol (PEG) generated predictable upshifts in migration through an SDS-PAGE gel. PEGylation of free thiols in the aggregated state (**Figure 5**) revealed that W42Q/C110S aggregates lost two free thiols per molecule, consistent with formation of one internal disulfide bond. The W42Q aggregates contained a major one-disulfide population and a minor, but still significant, population of molecules with two internal disulfide bonds. We infer that one of those bonds is 108-SS-110 and that its formation in the mutant is not required for aggregation.

**Figure 5:**
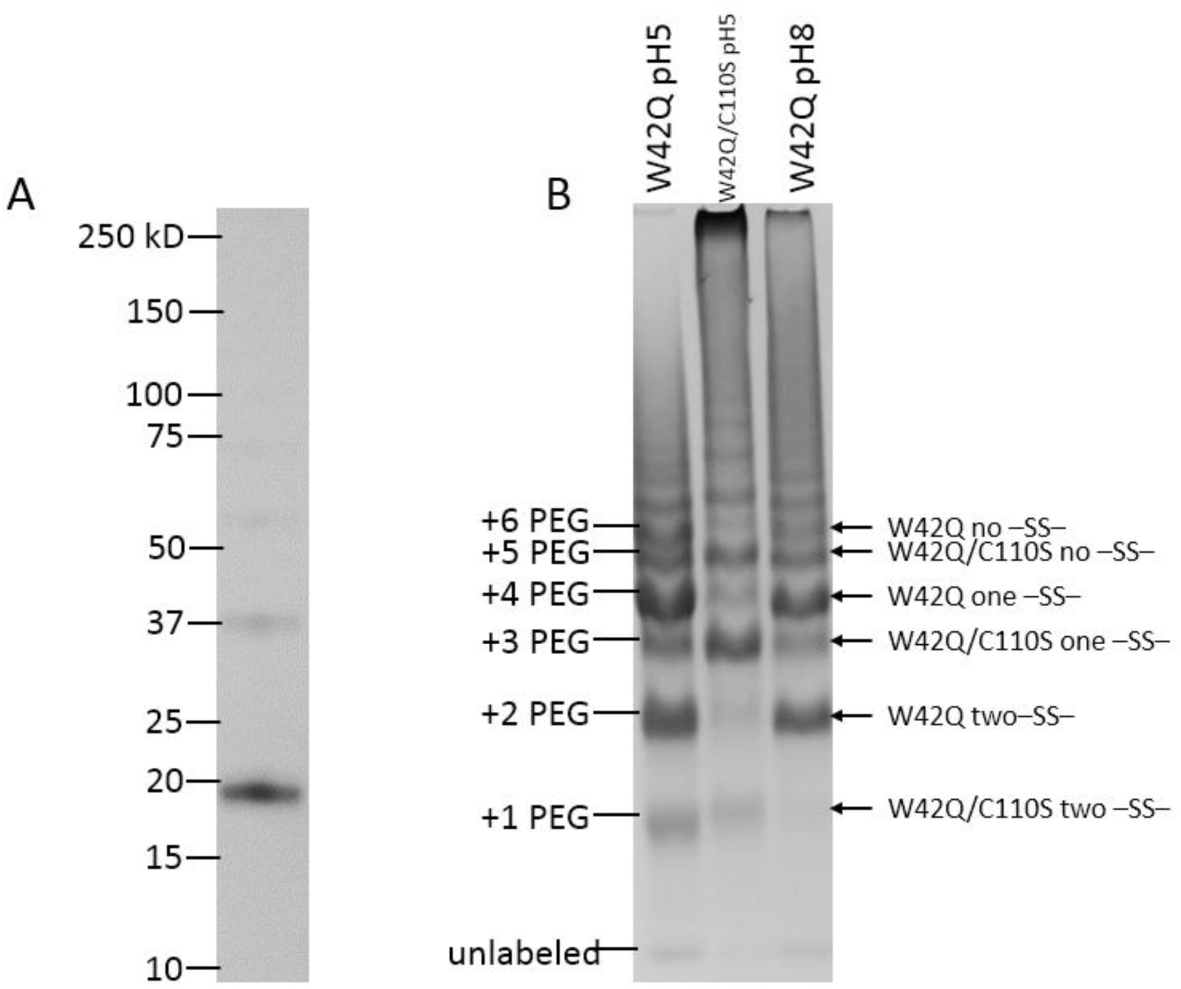
Distribution of disulfides in aggregated W42Q and W42Q/C110S by PEGylation. (A) The SDS-resolubilized aggregates of W42Q were predominantly monomeric by non-reducing SDS-PAGE, with only minor bands consistent with disulfide-crosslinked dimers and trimers visible on the gel. (B) Most aggregated monomers contained one internal disulfide, with a smaller fraction containing two, by PEGylation. *Left lane*: W42Q was mixed with oxidized WT, at increased concentration (75 μM of each protein) at pH 5.5, to generate aggregates while minimizing disulfide scrambling, and distributions of #free –SH groups per molecule in SDS-resolubilized aggregates were quantified by PEG-maleimide. The major band was found at +4 PEG (out of 6 Cys/molecule), indicating one internal disulfide per molecule, with a minor band at +2 PEG (two internal disulfides). *Middle lane*: The same was done for W42Q/C110S aggregates produced by mixing with oxidized C18T/C41A/C78A (75 μM of each protein) at pH 5.5. The triple-Cys mutant was used in order to ensure that any contamination from the non-W42Q proteins in the aggregates could be easily seen by SDS-PAGE upon PEGylation (as +1 or +3 bands in the W42Q lanes, and +4 or +6 bands in the W42Q/C110S lane, respectively). The predominant band for resolubilized monomers was at +3 PEG (out of 5 Cys/molecule), indicating one internal bond per molecule. Lack of a prominent minor band at +1 PEG indicated that monomers with two internal disulfides were extremely rare. *Right lane*: For comparison, the same was done for W42Q mixed with oxidized WT (40 μM of each protein) at pH 8, with results very similar to those obtained at pH 5.5. Since substantial numbers of monomers with two internal disulfides were found in W42Q aggregates, but not W42Q/C110S aggregates, we attribute the second disulfide to 108-SS-110.

### Oxidoreductase activity of HγD enables dynamic disulfide exchange among soluble HγD variants

The reversible C108-C110 disulfide of HγD bears similarity to the “C-X-C” motif of some thioredoxins and disulfide isomerases, as well as the redox-switchable chaperone Hsp33 (Derewenda et al., 2009; Jakob et al., 1999; Woycechowsky and Raines, 2003), so this gamma-crystallin may have a previously unrecognized oxidoreductase function. The Cys110 thiol group is solvent-accessible in the native structure (Basak et al., 2003), suggesting that disulfide interchange among HγD molecules may occur in solution. It was recently reported (Oka et al., 2015) that some protein disulfide isomerases are capable of dynamically exchanging disulfide bonds in solution, essentially forming a proteinaceous redox buffer in the endoplasmic reticulum. Given the high abundance of HγD in the lens core (up to ^~^10 mg/ml), it could potentially act as a supplementary redox buffer as glutathione levels are depleted with age.

To explore this possibility we incubated mixtures of HγD variants where one variant (or WT) started out as fully oxidized at the 108-110 site, and the other with those two residues fully reduced. The typical pKa of a Cys residue is ^~^8. Therefore, incubations were carried out either at pH 8 (permissive for thiol-disulfide interchange) or at pH 5 (inhibiting interchange). Disulfide transfer was determined by quantifying the distributions of the number of free thiols per molecule for each of the variants at the end of the incubation via the PEGylation/gel-shift assay As shown in **Figure 6**, when reduced WT protein was incubated with the oxidized triple-mutant C18T/C41A/C78A (abbreviated CCC) at pH 5, the vast majority of each respective polypeptide population was found at the migration positions expected for no disulfide exchange: the WT protein retained six free thiols per molecule (hence the prominent 6xPEG band), while the oxidized triple mutant had only one free thiol per molecule (hence the 1xPEG band). The band at 2xPEG was likely due to incomplete reaction between the CCC(ox) construct and the labeling reagent. The corresponding incubation mixture at pH 8 showed a significant decrease in the amount 6xPEG (WT) and 1xPEG (CCC(ox)) bands and a concomitant increase in the 3xPEG and 4xPEG bands, attributable to formation of reduced CCC mutant and oxidized WT, respectively.

**Figure 6:**
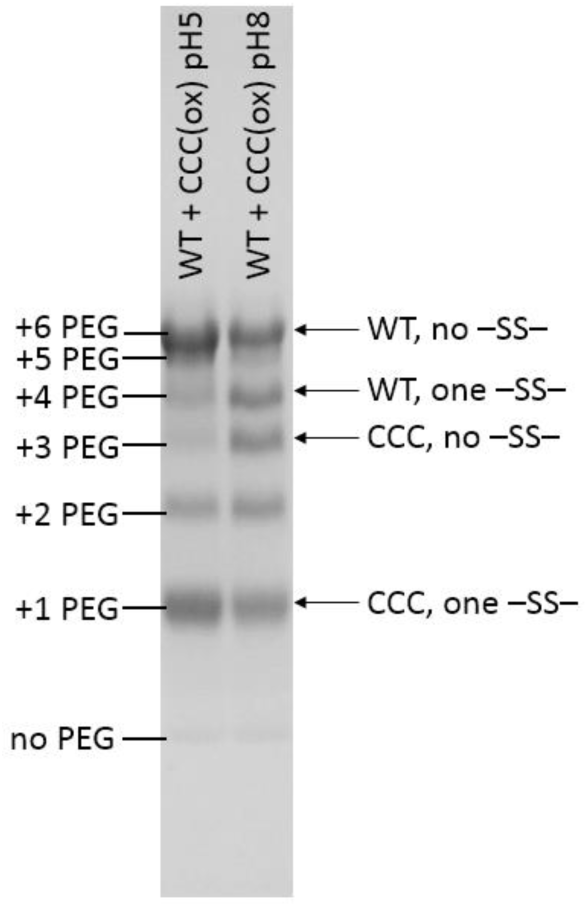
Evidence of disulfide transfer among soluble HγD molecules. An SDS-PAGE gel of samples reacted with maleimide-PEG to quantify the distribution of the number of free thiol groups per protein molecule is shown. When reduced WT protein was mixed with oxidized C18T/C41A/C78A triple mutant (*CCC(ox.)*) and incubated for 1 h at pH5, where disulfide exchange is inhibited (left lane), the resulting molecular population at the end of the reaction consisted largely of 6- and 1-free thiol populations, corresponding to the starting samples. When pH in the same mixture was set to 8, allowing disulfide exchange (right lane), after 1 h the population shifted toward 4- and 3-free thiol positions, corresponding to oxidized WT and reduced C18T/C41A/C78A, respectively.

In a parallel experiment, oxidized WT was incubated with reduced Y55A mutant – both constructs having the full complement of six Cys residues – and the result was analyzed by whole-protein electrospray mass spectrometry (**Figure SI 7**). As expected, the isotopically resolved molecular weight distribution of the WT population shifted upward as some of those molecules became reduced, which the mutant’s shifted downward, indicating oxidation. In a control experiment (**Figure SI 7**), no shifts were observed when neither variant had been oxidized, indicating that spontaneous formation of the 108-SS-110 bond was not a significant factor in these experiments.

### Formation of the 108-SS-110 disulfide conformationally strains the C-terminal domain

As we have previously demonstrated (Serebryany et al., 2016b), oxidative aggregation of W42Q/R mutants proceeds at temperatures ^~^10 °C below the onset of any detectable unfolding of the N-terminal domain. However, it is clear by inspection that formation of the 108-SS-110 disulfide should introduce strain into the native structure. Most thioredoxin-type motifs are “CXXC,” where XX are often Pro and Gly to enable the tight turn required to form the disulfide. In cases where “CXC” motifs are found, X is typically Gly (Derewenda et al., 2009; Woycechowsky and Raines, 2003) because its range of backbone angles is the broadest. We therefore hypothesized that the CSC motif found here was likely to be strained, which accounts for the ease of disulfide exchange yet may also contribute to conformational strain in the protein, potentially leading to partial unfolding.

To test this prediction experimentally, we studied both WT and the library of single and multiple Cys mutants in the WT background to determine whether 108-SS-110 formation indeed strained the native state. (The W42Q variant could not be oxidized efficiently due to extensive aggregation during the oxidizing treatment, as shown in **Figure SI 6**.) Differential scanning fluorometry experiments revealed that all variants capable of forming the 108-SS-110 bond had decreased apparent T_m_’s upon oxidation, relative to their reduced state (**Figure 7**). These decreases were not observed in constructs lacking either Cys108 or Cys110, but were present in all other cases. The conformational strain disappeared upon mild reduction, indicating that it was not due to any irreversible oxidative damage such as Met or Trp oxidation (**Figure SI 8**). The average downshift in DSF T_m_ attributable to the 108-SS-110 bond was 5.4 °C, with a standard deviation of 1.8 °C.

**Figure 7:**
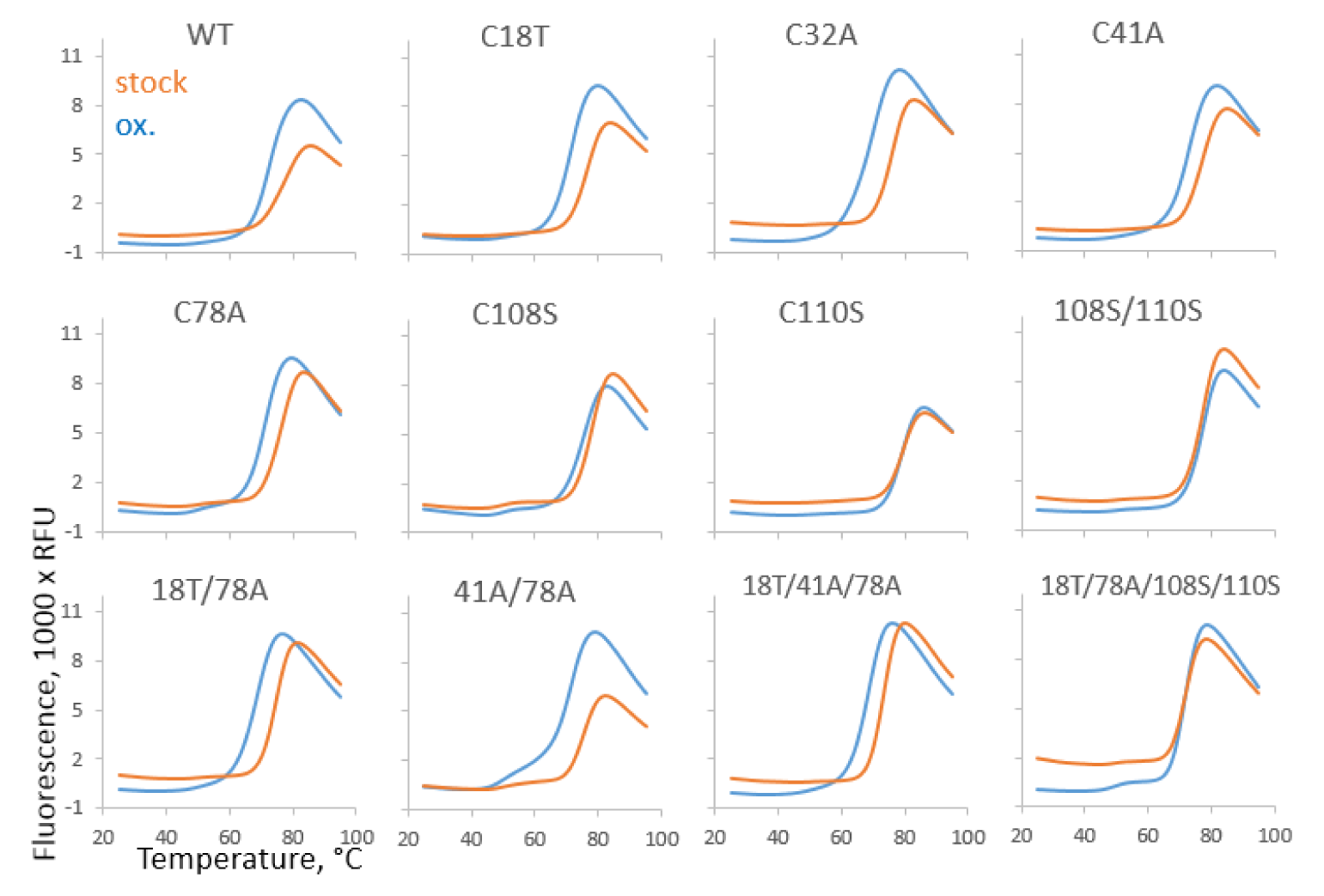
Destabilization due to formation of 108-SS-110. Differential scanning fluorometry traces for oxidized (orange) and reduced (blue) WT and a panel of Cys mutants, detected by fluorescence of the hydrophobicity probe SYPRO Orange. All the oxidized samples capable of forming the 108-SS-110 disulfide showed a shift of the melting transition to lower temperatures, indicating that presence of the disulfide bond was destabilizing. No such shifts were observed for C108S, C110S, C108S/C110S, or C18T/C78A/C108S/C110S – all mutants that are not capable of forming the 108-SS-110 disulfide. The shift is attributable to destabilization of the C-terminal domain upon oxidation (see Figure SI9). A shoulder at ^~^50 °C in the oxidized C41A/C78A double-mutant may be due to some misfolding or aggregation during the course of the measurement.

We next examined whether the conformational strain is global or confined to the C-terminal domain, which contains the disulfide. Differential scanning fluorometry assumes that the reporter fluorophore – in this case SYPRO Orange – binds to all relevant molten-globule states. However, DSF measurements conducted on the W42Q mutant (**Figure SI 9**) revealed only one melting transition, rather than the two transitions previously reported for this mutant by calorimetry, guanidinium denaturation, and intrinsic fluorescence (Ji et al., 2013; Serebryany and King, 2015; Serebryany et al., 2016a; Serebryany et al., 2016b). The temperature of the transition was above the previously measured T_m_ of the N-terminal domain of the W42Q protein and close to its C-terminal T_m_. Thus, SYPRO Orange likely bound preferentially to the molten-globule state of the C-terminal domain rather than the N-terminal domain. This surprising observation may help explain why chaperones fail to recognize the partial unfolding of the N-terminal domain in aggregation-prone, cataract-associated variants of this protein (Moreau and King, 2012a; Serebryany et al., 2016a). Apparently, unfolding of the N-terminal domain did not form a sufficiently hydrophobic binding pocket. This may be due in part to some of the N-terminal domain’s residues being sequestered upon partial unfolding by interaction with the C-terminal domain, for example as proposed in (Serebryany et al., 2016b).

To confirm the effect of the disulfide strain by a complementary label-free measurement, we carried out differential scanning calorimetry (DSC) experiments at pH5, to inhibit any disulfide exchange, comparing stock, oxidized, and oxidized-then-reduced WT HγD, as well as identically treated C18T/C41A/C78A triple-mutant. The results are summarized in **Table 1**. The stock WT protein was destabilized only modestly (3-4 °C) at lower pH, compared to the previous neutral-pH measurements (Serebryany et al., 2016b). However, oxidation led to a downshift of ^~^10 °C in the melting transition of one domain in the WT and ^~^7 °C in the triple-mutant, while the other melting transition remained unchanged in both cases (**Figure SI 10**). We attribute the redox-dependent melting transition to the C-terminal domain, since the redox-active Cys pair is located there. The fitted calorimetric enthalpies for the two domains (shown in **Table 1**) were consistent with this assignment. Although these calculated enthalpies may be subject to variations in protein concentration or to aggregation during the course of the experiment, the overall pattern was clear: enthalpy of the C-terminal domain was significantly higher than that of the N-terminal domain in reduced samples, but dropped to become nearly identical to it following 108-SS-110 formation. The oxidized-then-reduced samples yielded traces and melting transitions very similar to those of the never-oxidized samples, indicating that destabilization due to oxidation was almost entirely redox-reversible and thus attributable to conformational strain induced by the 108-SS-110 disulfide.

**Table 1:**
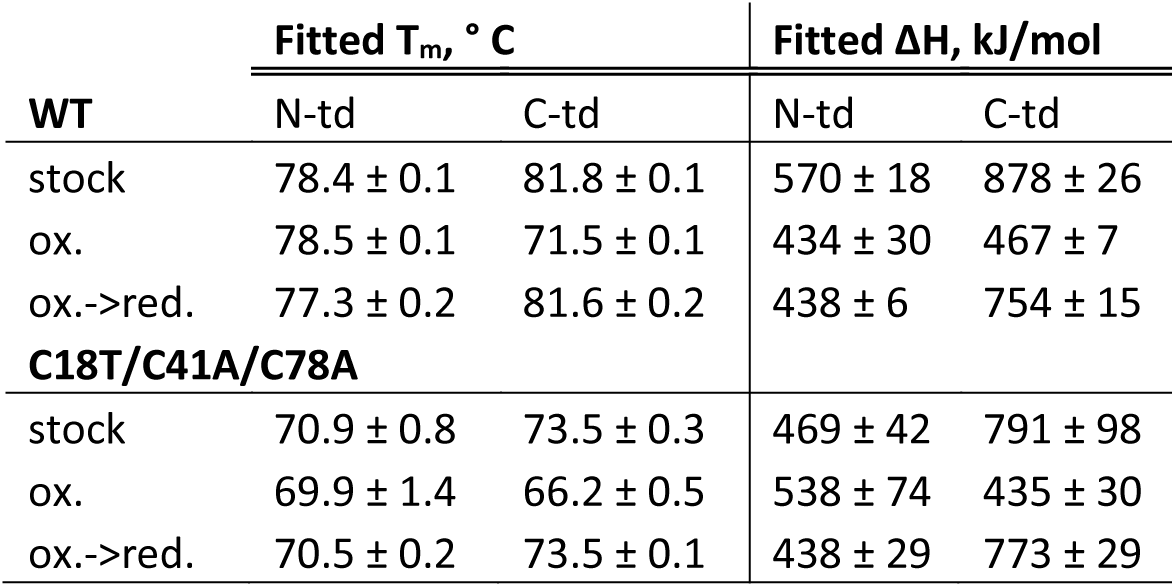
Melting temperatures and enthalpies from fitting of WT and C18T/C41A/C78A HγD melting traces as a function of redox treatment, measured by differential scanning calorimetry at pH 5, reported as the average ± 2^∗^S.E.M. of three replicates. N-td = N-terminal domain; C-td = C-terminal domain.

| | Fitted $T_m$, °C | | Fitted $\Delta H$, kJ/mol | |
| --- | --- | --- | --- | --- |
|  | N-td | C-td | N-td | C-td |
| **WT** |  |  |  |  |
| stock | 78.4 ± 0.1 | 81.8 ± 0.1 | 570 ± 18 | 878 ± 26 |
| ox. | 78.5 ± 0.1 | 71.5 ± 0.1 | 434 ± 30 | 467 ± 7 |
| ox.->red. | 77.3 ± 0.2 | 81.6 ± 0.2 | 438 ± 6 | 754 ± 15 |
| **C18T/C41A/C78A** |  |  |  |  |
| stock | 70.9 ± 0.8 | 73.5 ± 0.3 | 469 ± 42 | 791 ± 98 |
| ox. | 69.9 ± 1.4 | 66.2 ± 0.5 | 538 ± 74 | 435 ± 30 |
| ox.->red. | 70.5 ± 0.2 | 73.5 ± 0.1 | 438 ± 29 | 773 ± 29 |

The W130E oxidation-mimicking mutation in the C-terminal domain of HγD opens up a distinct non-native, non-amyloid aggregation pathway, which is chaperone-suppressible yet also partially redox-dependent (Serebryany et al., 2016a). It is likely that the destabilization produced by the C-terminal mutation is in synergy with the conformational strain produced by the 108-SS-110 bond, and thus the disulfide indirectly promotes aggregation of this cataract-associated protein. Notably, UV irradiation, one of the best-characterized causes of cataract, is known to damage Trp130 in HγD (Moran et al., 2013), and the UV-induced aggregation pathway requires oxidation (Schafheimer and King, 2013). The N-terminal domain of HγD derives part of its stability from its interface with the C-terminal domain; disrupting this interface leads to N-terminal destabilization (Flaugh et al., 2005a, b; Mills et al., 2007; Moreau and King, 2009). The earliest detectable unfolding of the W42Q protein occurs at ^~^43 °C (Serebryany and King, 2015; Serebryany et al., 2016b), so if the 108-SS-110 bond forms in that variant, and the resulting destabilization propagates to the domain interface, it may be sufficient to increase the population of aggregation-prone intermediates near 37 °C and in this way allosterically promote formation of the non-native 32-SS-41 disulfide bond in the N-terminal domain that is required for aggregation. We have not found evidence of such an allosteric effect; however, a subtle allosteric effect on the N-terminal domain’s dynamics near physiological temperature cannot be ruled out.

## Discussion

This work began with the mystery of how the kinetically and thermodynamically stable (Flaugh et al., 2006; Kosinski-Collins and King, 2003) WT HγD is able to catalyze aggregation of its misfolding-prone W42Q variant without itself aggregating. This behavior is the inverse of the classical prion-like aggregation model, where misfolding-prone variants catalyze misfolding and aggregation of natively folded ones. However, the key to this “inverse prion” paradox turned out to be redox chemistry, rather than conformational templating. The W42Q mutation, by partially destabilizing the N-terminal domain, enables two Cys residues that are distant and buried in the natively folded N-terminal domain to become solvent-exposed and transiently approach each other (Serebryany et al., 2016b). We have found that a redox-active disulfide in the C-terminal domain of the WT protein can then be transferred to these newly exposed N-terminal Cys residues in the mutant. The products of this bimolecular redox reaction are a soluble, fully reduced WT molecule and a W42Q molecule bearing a non-native internal disulfide that locks its N-terminal domain in an aggregation-prone intermediate state (**Figure 8**). The structure of this intermediate, and of the resulting aggregated state, has been proposed (Serebryany et al., 2016b). The intermediate structure – detachment of the N-terminal β-hairpin – is consistent with that derived from single-molecule force spectroscopy (Garcia-Manyes et al., 2016), and evidence of the specific disulfide bond that traps this intermediate has been found in patient lenses and correlates strongly with age-onset cataract (Fan et al., 2015). It is worth noting that the latter two studies were carried out on un-mutated proteins, and that a large variety of congenital cataract-linked point mutations in the γ-crystallins (including the W42R mutation) cluster near this N-terminal hairpin (Serebryany and King, 2014). These observations raise the possibility that the W42Q mutation merely increases the population of a conformational intermediate that is already accessible, albeit extremely rare, in the wild-type protein, and that other mutations and age-related post-translational modifications converge on the same aggregation-prone intermediate. In this model, the fact that the oxidized WT HγD does not aggregate on experimentally measurable timescales is simply due to high kinetic stability of its N-terminal domain.

**Figure 8:**
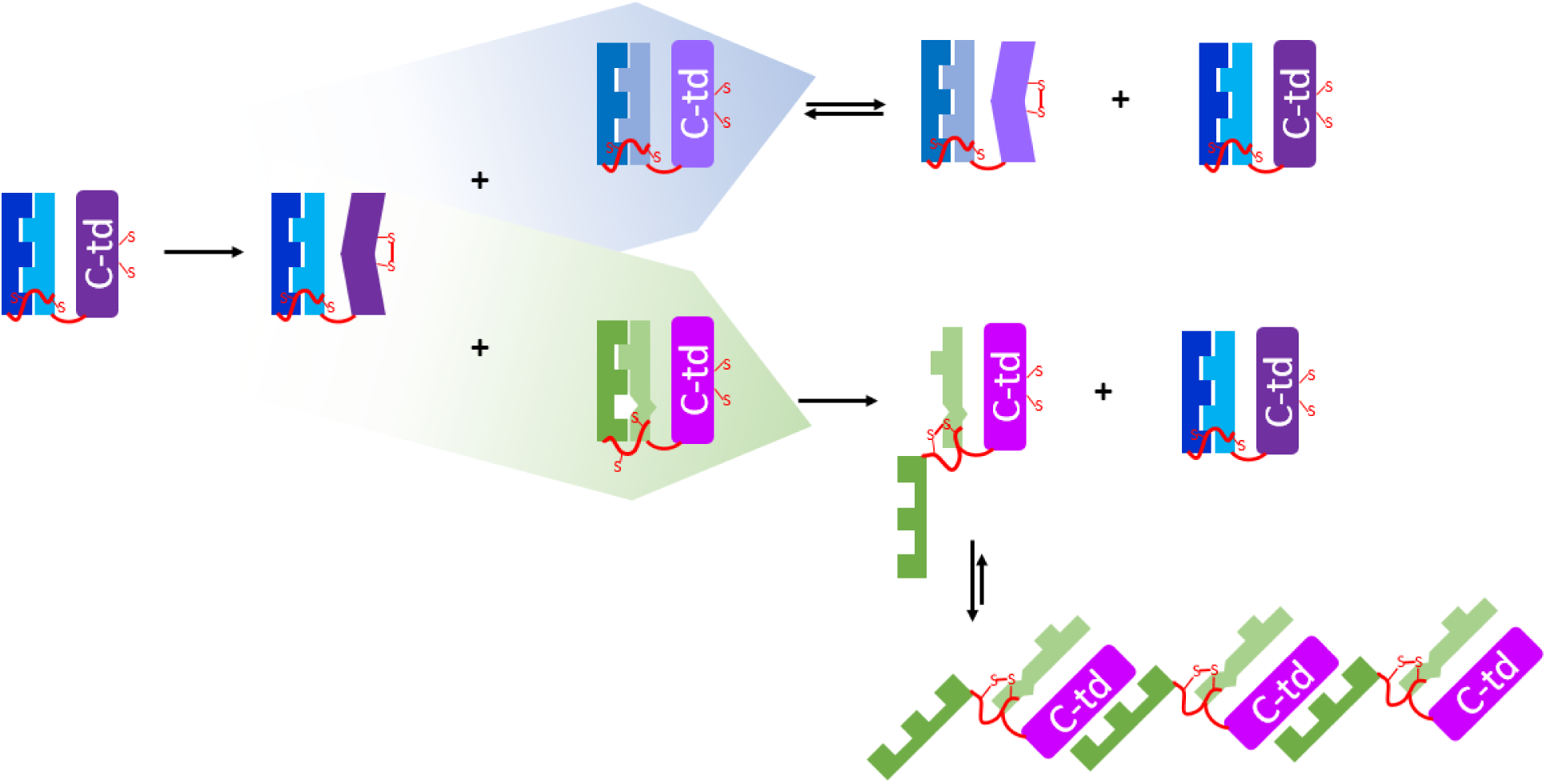
An illustration of the “redox hot potato” model of inverse-prion aggregation mediated by dynamic disulfide bonds. The WT, undamaged protein (*left*) may become oxidized, forming a conformationally strained and hence redox-active disulfide. This disulfide can then migrate to either (*blue arrow*) a reduced WT or WT-like protein, in a fully reversible reaction, or (*green arrow*) a damaged or mutated protein (here indicated by a perturbed native interaction in the N-terminal domain). In the latter case, it may migrate to a different location in the molecule, forming a conformationally relaxed and hence redox-inactive disulfide between Cys residues that are normally buried in the native state. This non-native disulfide then traps a partially unfolded intermediate in a reaction that is essentially irreversible under mildly oxidizing conditions. The intermediate cannot revert to the native state and therefore partitions to aggregated structures. We have previously proposed a structural model for this aggregated state in (Serebryany et al., 2016b).

The specific model proposed here postulates transfer of the Cys108-Cys110 disulfide from WT HγD to a non-native N-terminal disulfide (Cys32-Cys41) of the W42Q mutant. Of the six Cys residues in HγD, five are well conserved among the γ-crystallins; Cys110 is the exception, being present in humans and chimpanzees, but not in chicken, mouse, rat, or bovine γD sequences (Kong and King, 2011). Remarkably, however, each of those species has a Cys residue in position 110 in the closely related γC-crystallin, whereas human and chimpanzee versions of γC have Ser110 instead. Thermodynamically, HγD is more stable than HγC (Serebryany and King, 2014), and human lens proteomics has revealed that HγD persists longer than HγC, at least in its soluble form, becoming the most abundant soluble γ-crystallin by age ^~^55 (Ma et al., 1998). Thus, the apparent evolutionary switch of the Cys108-Cys110 pair from γC to γD might be a consequence of increased lifespan of the organism. At the same time, it is worth noting that at least one study identified other sequence-proximal disulfides in the γ-crystallins: the Cys78-Cys79 bond in HγC and the Cys22-Cys24 bond in HγS (Hanson et al., 1998). It is not yet known whether these bonds are dynamic. Conversely, previous research also identified other non-native internal disulfides in cataractous lenses, particularly in β-crystallins (Takemoto, 1997a, b), as well as one such bond in the lens-specific chaperone αA-crystallin, which appeared to diminish its chaperone activity as well as contribute to aggregation (Cherian-Shaw et al., 1999; Takemoto, 1996). More recently many disulfide bonds were discovered in non-crystallin proteins in the lens (Wang et al., 2017). It is possible that many of these long-lived lens proteins fall within the redox “hot potato” model we have proposed.

What, then, is the possible fitness benefit of these sequence-proximal and likely redox-active disulfides? A possible explanation is that glutathione levels become depleted in lens tissue during the course of aging (Friedburg and Manthey, 1973), and the lens core gradually becomes impermeable even to glutathione generated in the lens cortex or present in vitreous humor (Sweeney and Truscott, 1998), resulting in increased disulfide formation in lens cytoplasmic proteins (Wang et al., 2017). The γ-crystallins have always been thought of as purely structural proteins with no known biochemical function aside from their structural stability and optical properties (Bloemendal et al., 2004; Moreau and King, 2012b; Serebryany and King, 2014). The search for the inverse-prion mechanism led us to a previously unrecognized oxidoreductase activity in wild-type HγD. Dynamic exchange of disulfide bonds among the protein molecules makes reduced and oxidized forms of HγD a redox couple. Given the high abundance of γ-crystallins, we speculate that they may constitute a protein-based redox buffer in the aging, glutathione-depleted lens. If so, there is some evidence that this buffering capacity is enzymatically regulated *in vivo*. Early proteomic studies of the human eye lens indicated that Cys110 of HγD is partially (^~^30-70%) methylated (Lapko et al., 2003). The significance of this observation was not known at the time, but can now be interpreted as a likely regulation mechanism for the protein’s oxidoreductase activity. Like Cys110 of HγD, Cys79 of HγC and Cys24 of HγS have also been found to be partially methylated *in vivo* (Lapko et al., 2002, 2003), suggesting a common physiological regulation mechanism. Cysteine methylation is an enzyme-catalyzed reaction *in vivo* (Clarke, 2013). Our biochemical observations thus open up the possibility that an enzymatic pathway in the lens core actively regulates the redox buffering capacity of the crystallin proteome and may influence the age of onset of cataract.

Moreover, we have revealed a likely failure mode of such a protein-based redox homeostasis system: while sequence-proximal disulfides are kinetically favorable, non-native ones are favored thermodynamically due to their propensity to aggregate. Destabilizing mutations or modifications of the HγD core, such as oxidation of Trp residues, may generate disulfide sinks analogous to the W42Q mutant. In general, we propose a redox “hot potato” mechanism (**Figure 9**) in which polypeptides with lower kinetic stability (due to mutations or post-translational modifications) become trapped in aggregation-prone intermediate states and insolubilized as a consequence of accepting disulfides from more stable variants. The result is, essentially, a kinetic stability competition among protein variants competent for disulfide exchange, with the conformational “weakest link” variant driven into an aggregated state upon receiving a disulfide. To participate in such a competition, the main features of the protein should be: (1) ability to exchange disulfides – or perhaps other reversible modifications – under physiologically relevant conditions; and (2) a folding landscape that contains aggregation-prone intermediates that become more populated upon receiving the modification – in this case, a disulfide bond – effectively serving as a sink for modified proteins. This process may have played a role in the evolution of the exceptional stability of the γ-crystallins or other long-lived proteins.

**Figure 9:**
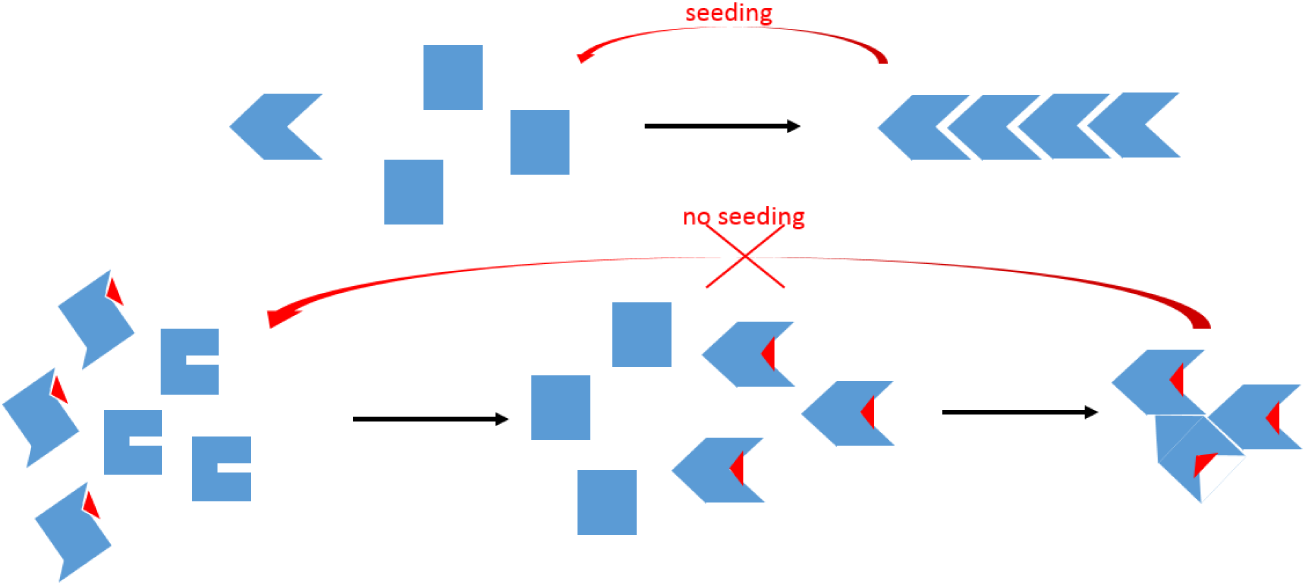
Schematic representation of a classical prion and a “hot potato” inverse prion. The classical prion-like aggregation model (*top*) involves conformational templating of misfolded states onto native states. One consequence is the well-known seeding effect, whereby mixing aggregates or fragments of aggregates with the starting solution dramatically accelerates aggregation. The proposed “hot potato” model (*bottom*) involves interaction between a native-like high-energy state trapped by a post-translational modification and a conformationally labile variant (generated by mutation or a different post-translational modification), leading to regeneration of the fully native state in the former at the cost of generating the aggregation-prone misfolded state in the latter. Notably, since conformational templating does not play a primary role in the aggregation reaction, no seeding effect need be expected, and indeed no significant seeding is observed (Serebryany et al., 2016b).

## Methods

*Site*-*directed mutagenesis* was carried out using either the Quikchange II kit (Agilent, Santa Clara, CA) or the Q5 kit (New England Biolabs, Ipswich, MA) according to the manufacturers’ instructions. Resulting plasmids, amplified in the XL-1 *E. coli* strain (Agilent), were confirmed by sequencing and transformed into the BL21-RIL strain of *E. coli* for expression.

*Protein expression and purification* was carried out largely as described (Serebryany and King, 2015; Serebryany et al., 2016b) with some modifications. Briefly, overnight starter cultures of BL21-RIL *E. coli* were inoculated in 10-40 ml reconstituted SuperBroth media (Teknova, Hollister, CA) supplemented with ampicillin and chloramphenicol. Expression cultures using the same media in standard 2L flasks were inoculated at 1:100 from the starters and grown at 37 °C with shaking at 250 rpm for 6-8 h until reaching stationary phase (OD_600_ ^~^ 2). Cultures were then chilled to 18 °C for 30 min and induced overnight at 18 °C by adding IPTG (Promega, Madison, WI) to 1 mM final concentration. Harvested cells were resuspended in the presence of protease inhibitors (Complete-mini, EDTA-free, Roche USA, Branford, CT) and stored at −70 °C until used.

Following lysis by sonication in the presence of DNase and lysozyme, purification was carried out by salting out with ammonium sulfate as described (Serebryany and King, 2015). The 50% ammonium sulfate supernatant was dialyzed at 4 °C overnight against 4 L of sample buffer (10 mM ammonium acetate, 50 mM NaCl) to remove the ammonium sulfate. The dialysis step typically caused significant precipitation, attributable to peptidoglycan contaminants. Following centrifugation, these samples were then further scrubbed of lipids and peptidoglycans by passage through an anion-exchange column comprised of two 5 ml Q-Sepharose columns (GE Healthcare, Marlborough, MA) in tandem equilibrated in sample buffer. Crystallins eluted in the flow-through. The ion-exchanged samples were concentrated using VivaSpin Turbo 5K MWCO centrifugal filters (Sartorius, Goettingen, Germany), loaded onto a Superdex75 26×600 column (GE Healthcare), and separated at 2-3 ml/min flow rate at room temperature. Resulting fractions were collected and stored at 4 °C. SDS-PAGE (Criterion, BioRad, Hercules, CA) was used to determine >95% purity of the samples. Proteins with the W42Q mutation were often found to have a minor degradation product at ^~^10 kD in the gel, but its presence did not noticeably affect the overall aggregation behavior. Proteins were concentrated for storage at 4 °C to 100-500 μM; concentrations were determined A_280_ using NanoDrop 2000 instrument (Thermo Fisher, Waltham, MA).

*In-vitro oxidation* to form the 108-SS-110 disulfide bond was carried out by mixing 20 μM CuSO4 and 60 μM phenanthroline (MilliporeSigma, Burlington, MA) in sample buffer at room temperature, to chelate the copper, and then adding the protein to 20 μM final concentration. This mixture, typically in 5-10 ml volume, was incubated at room temperature for 1-2 h in closed 15-ml conical tubes, followed by addition of 1 mM EDTA and three rounds of dialysis through metal-free SpectraPor 7 dialysis membrane (Spectrum Labs, Breda, The Netherlands) at 4 °C against sample buffer for at least 3 h at a time (at least once overnight). The first round of dialysis included 1 mM EDTA in the sample buffer to fully chelate and remove the Cu^2+^ ions from the sample and avoid any copper-induced aggregation or oxidation during the subsequent experiments (Quintanar et al., 2016). To ensure that biochemical differences between the oxidized and reduced samples were due to reversible oxidation, i.e., disulfide formation, oxidized samples were subjected to mild reducing treatment – 1 mM dithiothreitol for 2 h at 37 °C – generating the oxidized->reduced (ox.->red.) controls.

*Aggregation assays* were carried out at 37 °C unless otherwise indicated, using half-area clear polypropylene plates (Greiner Bio-One North America, Monroe, NC) in a PowerWave HT plate reader (BioTek, Winooski, VT) in 100 μl volume without shaking. ^~^20% of the sample typically evaporated during the course of the 3 h aggregation experiment. Resulting path length was approximately 0.5 cm, so the reported turbidity values are expected to be ^~^50% lower than in the typical 1-cm path length spectrophotometer cuvette. Aggregation occurred on a comparable time scale in other reaction vessels, such as capped microcentrifuge or thermocycler tubes.

*Differential Scanning Fluorometry* was carried out using 1:1 mixture of sample buffer and sodium phosphate (pH7, 100 mM) buffer in order to maintain neutral pH during heating, in 96-well format in a CFX96 RT-PCR instrument (BioRad). 1X SYPRO Orange dye (Life Technologies, Carlsbad, CA) was added as the hydrophobicity probe; control (no-protein) samples were subtracted. Temperature ramp was at 1 °C / min between 25 °C and 95 °C. Melting temperatures were defined as the minima (rounded to the nearest ° C) of the derivative of the empirically fit sigmoid functions in the CFX Manager software (BioRad).

*Differential Scanning Calorimetry* was carried out as reported previously (Serebryany et al., 2016b), except in pH5 buffer (20 mM ammonium acetate, 50 mM NaCl) to inhibit disulfide exchange during the course of the measurement. Proteins were at 25 μM concentration. Samples were kept at 4 °C until immediately prior to analysis, which ran from 25 to 95 °C. Each melting trace was fitted to a sum of two 2-state scaled models using the NanoAnalyze software (TA Instruments, New Castle, DE).

*PEGylation assays* were carried out as follows. Samples were denatured for 5 min at 95 °C in pH5 buffer (50 mM ammonium acetate) with 5% w/v SDS at [protein] of ^~^30 μM in a total volume of 10 μl per sample. Once cooled to room temperature, 10 μl of pH8 buffer (100 mM sodium phosphate) was added to neutralize, followed by 6 μl of 1 mM maleimide-conjugated polyethylene glycol, M_r_ ^~^5000 (MilliporeSigma) and 4 μl of 4X NuPAGE SDS-PAGE gel-loading buffer (Thermo Fisher). The reaction mixtures were incubated at 50 °C for 2 h, then analyzed directly by SDS-PAGE with Coomassie stain (Thermo Fisher).

*Isotopically resolved intact mass determination* was accomplished by electrospray ionization mass spectrometry. The protein samples were analyzed on a Bruker Impact II q-TOF mass spectrometer equipped with an Agilent 1290 HPLC. The separation and desalting was performed on an Agilent PLRP-S Column (1000A, 4.6 × 50 mm, 5 μm). Mobile phase A was 0.1% formic acid in water and mobile phase B was acetonitrile with 0.1% formic acid. A constant flow rate of 0.300 ml/min was used. Ten microliters of the protein solution was injected and washed on the column for the first 2 minutes at 0%B, diverting non-retained materials to waste. The protein was then eluted using a linear gradient from 0%B to 100%B over 8 minutes. The mobile phase composition was maintained at 100%B for 1 minutes and then returned to 0%B over 0.1 minute. The column was re-equilibrated to 0%B for the next 5.9 minutes. A plug of sodium formate was introduced at the end of the run, to perform internal m/z calibration to obtain accurate m/z values. The data were analyzed using Bruker Compass DataAnalysis^™^ software (Version 4.3, Build 110.102.1532, 64 bit). The charge state distribution for the protein produced by electrospray ionization was deconvoluted to neutral charge state using DataAnalysis implementation of the Maximum Entropy algorithm. Predicted isotope patterns were calculated at the resolving power of 50,000 and compared with isotopically resolved neutral mass spectra calculated using Maximum Entropy from the experimental charge state distribution.

*Disulfide mapping* was carried out using LC-MS/MS on an Orbitrap Fusion Lumos Tribrid Mass Spectrometer (Thermo Fisher) equipped with EASY nanoLC 1000 pump (Thermo Fisher). Peptides were separated onto a 100 μm inner diameter microcapillary Kasil frit trapping column packed first with approximately 5 cm of C18 Reprosil resin (5 μm, 100 Å, Dr. Maisch GmbH, Ammerbuch-Entringen, Germany). Separation was achieved through applying a gradient from 5–27% acetonitrile in 0.1% formic acid over 90 min at 200 nl/min. Electrospray ionization was enabled through applying a voltage of 2 kV using a home-made electrode junction at the end of a Thermo Nano Viper 1200 75um × 550mm column. The LUMOS Fusion was operated in the data-dependent mode for the mass spectrometry methods. The mass spectrometry survey scan was performed in the range of 395 –1,800 m/z at a resolution of 6 × 10^4^, with HCD fragmentation in the Fusion trap using a precursor isolation width window of 2 m/z. HCD Collision energy was set at 32 volts; isolation window was 3 Da with 30k Orbitrap; resolution for MS2 EThCD scan was 200 milliseconds; activation time was 10 ms; AGC was set to 50,000. Ions in a 10 ppm m/z window around ions selected for MS2 were excluded from further selection for fragmentation for 60 s.

WT(ox.) samples were treated with either chymotrypsin alone or the combination of trypsin and chymotrypsin. Samples were not carbamidomethylated or reduced; rather, the pH of the digestion mixture was kept neutral or below at all times so as to minimize disulfide scrambling, as in our previous report (Serebryany et al., 2016b). W42Q aggregates were pelleted by centrifugation at 12,000 × g for 5-10 min, rinsed with 3 ml of sample buffer, and then resuspended in pH5 ammonium acetate buffer. The suspension was centrifuged again; the supernatant was collected as the “reversible” fraction, while the remaining solid material was considered “irreversible.” Both were then denatured in SDS (the irreversible fraction could not be fully solubilized even in 5% SDS at 80 °C) and trypsinized using the S-trap method (Protifi, Huntington, NY), with samples collected at 15, 45, and 90 minutes of incubation. Results were analyzed using Comet software, and additionally verified by probability scores using PeptideProphet software, with searches against the relevant subset of the cRAP database of common contaminants (Mellacheruvu et al., 2013), such as human keratins, within the Trans-Proteomic Pipeline. Only peptides with probability scores >0.95 were included in the analysis.

Top-down proteomics were used to confirm disulfide assignments, using the same equipment as described above. WT(ox.) sample was co-digested with chymotrypsin and GluC under incomplete-digestion conditions. Fragment MS1 spectra were deconvoluted by Maximum Entropy and assigned on the basis of their molecular weight, as well as the weights of sub-fragments arising from in-source decay.

## Supplementary Information

**Table SI1:**
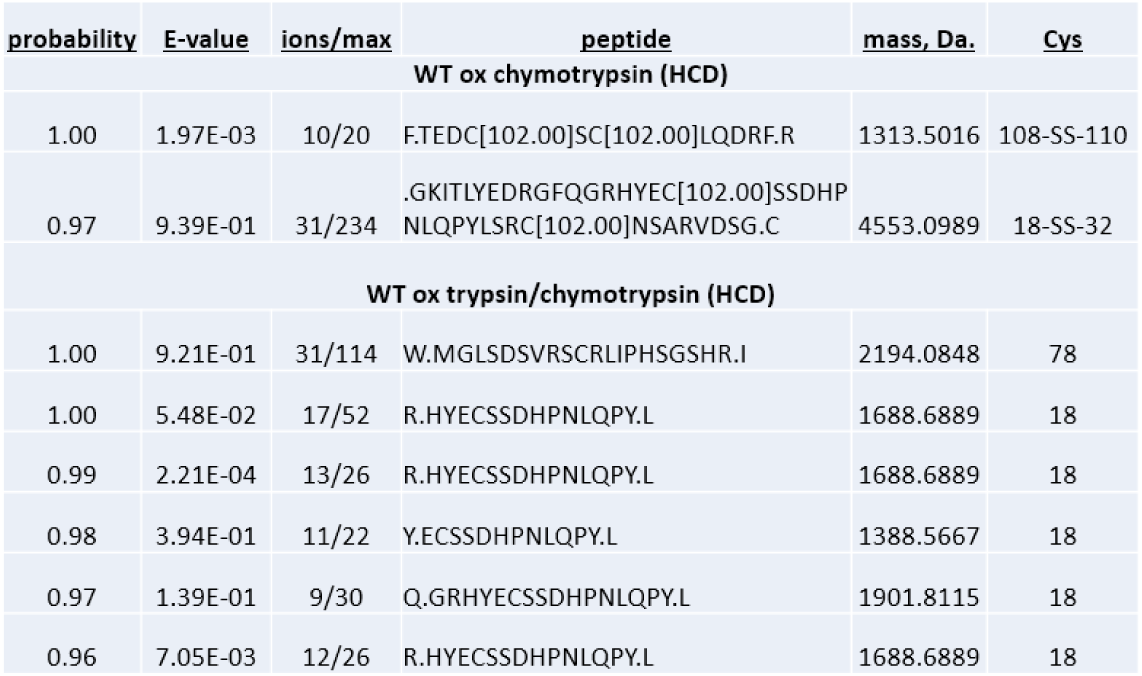
mass spectrometric identification of the 108-SS-110 bond. Oxidized WT was denatured for 5-10 min at 80 °C and then digested with either chymotrypsin alone or a mixture of trypsin and chymotrypsin. Comet searches on the resulting LC/MS/MS spectra were carried out including a −1.007 Da. variable modification on the Cys (loss of one proton). Peptide hits whose masses corresponded to only one such modification were discarded as false positives. Peptides identified with a PeptideProphet probability score of at least 0.95 are shown. A strong identification was made for the 108-SS-110 bond, with a less-confident signal for 18-SS-32, in the oxidized WT sample; on the other hand, high-confidence hits showing reduced forms of Cys18 and Cys78 were also found. Cys41 could not be detected in these experiments, likely due to inefficient digestion or poor ionization.

**Figure SI2:**
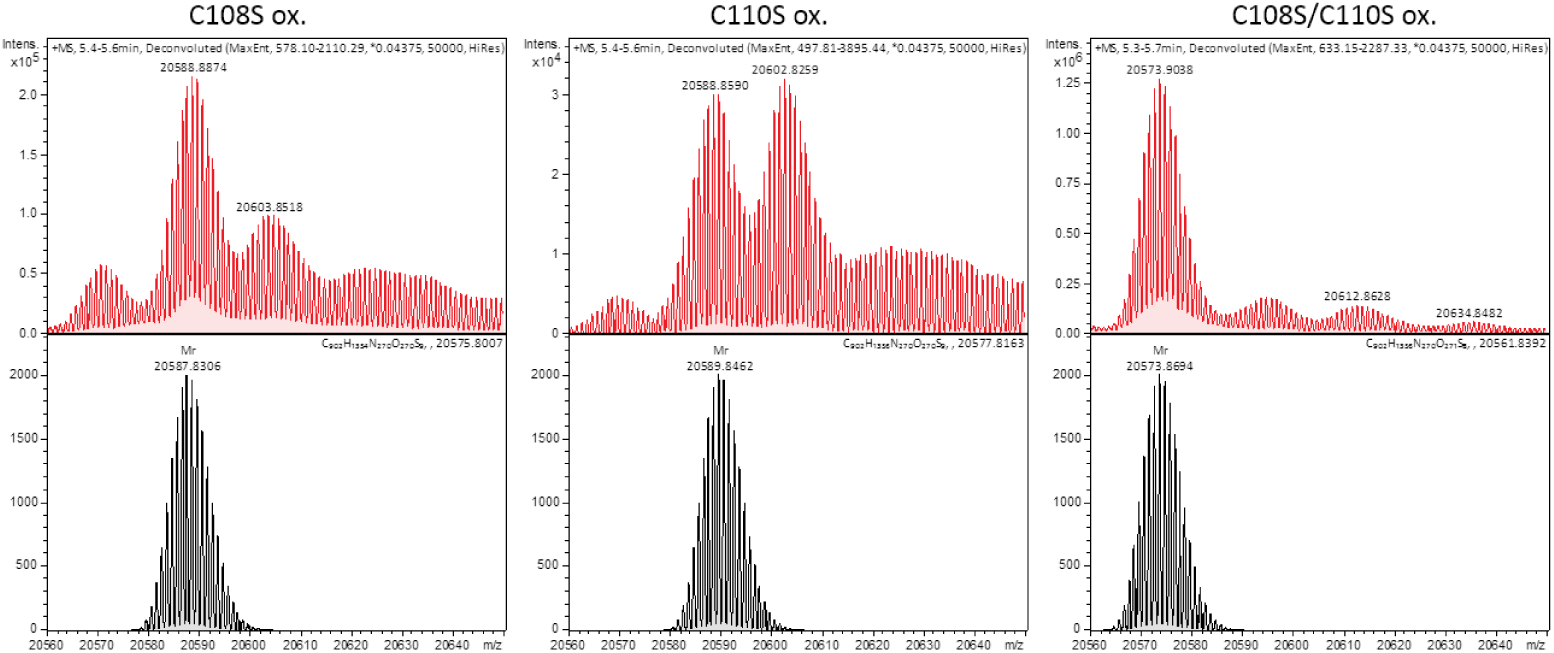
108-SS-110 formation may protect against irreversible side chain oxidation. Whole-protein isotopically resolved mass spectra (red) and predicted spectra (black) indicate that no internal disulfide forms upon copper/phenanthroline treatment when the protein lacks Cys108 or Cys110 (no −2Da shift). However, significant amounts of a +16Da. product are observed, particularly in the case of the oxidized C110S mutant, but to a lesser extent also C108S, which suggests the presence of an oxygen adduct. The most likely interpretation is oxidation of a Met residue. Remarkably, this oxidation does not occur when both Cys108 and Cys110 are absent. We hypothesize that it may arise when an unstable Cys oxidation product, such as sulfenic acid, is resolved to a methionine sulfoxide.

**Table SI3:**
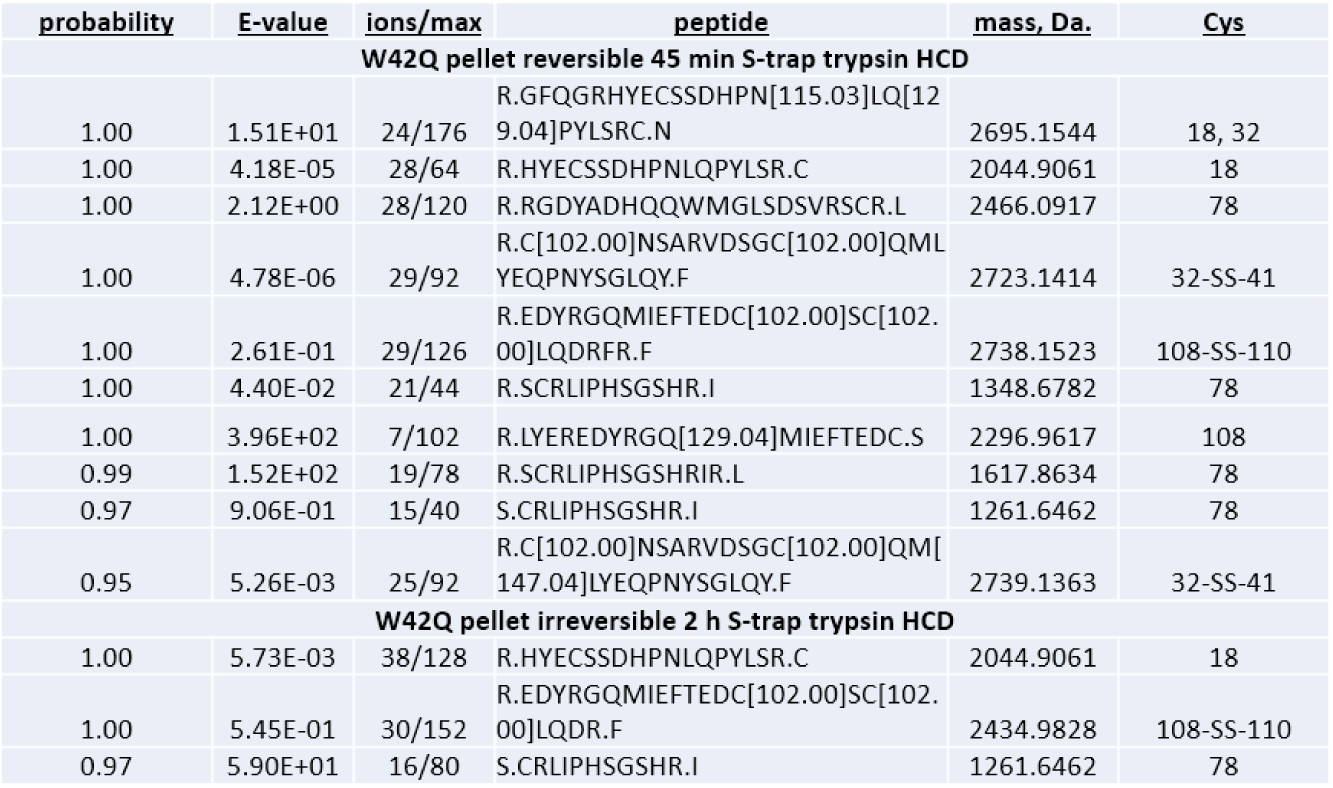
Mass spectrometric identification of prevalent disulfides in W42Q aggregates. W42Q aggregates were divided into “reversible” and “irreversible” based on the ease of their resolubilization in fresh pH5 buffer. Both were digested with trypsin in the presence of SDS as denaturant, using the S-trap procedure. Comet searches on the resulting LC/MS/MS spectra were carried out including a −1.007 Da. variable modification on the Cys (loss of one proton). Peptide hits whose masses corresponded to only one such modification were discarded as false positives. High-confidence hits include identification of the 32-SS-41 disulfide, as well as a less-confident identification of 108-SS-110. By contrast, Cys18 and Cys78 are found exclusively in their reduced forms. One peptide was identified as having both Cys18 and Cys32, reduced, but the presence of two apparent deamidations simultaneously (indicated by N[115.03] and Q[129.04]) suggests this peptide is rare.

**Figure SI 4:**
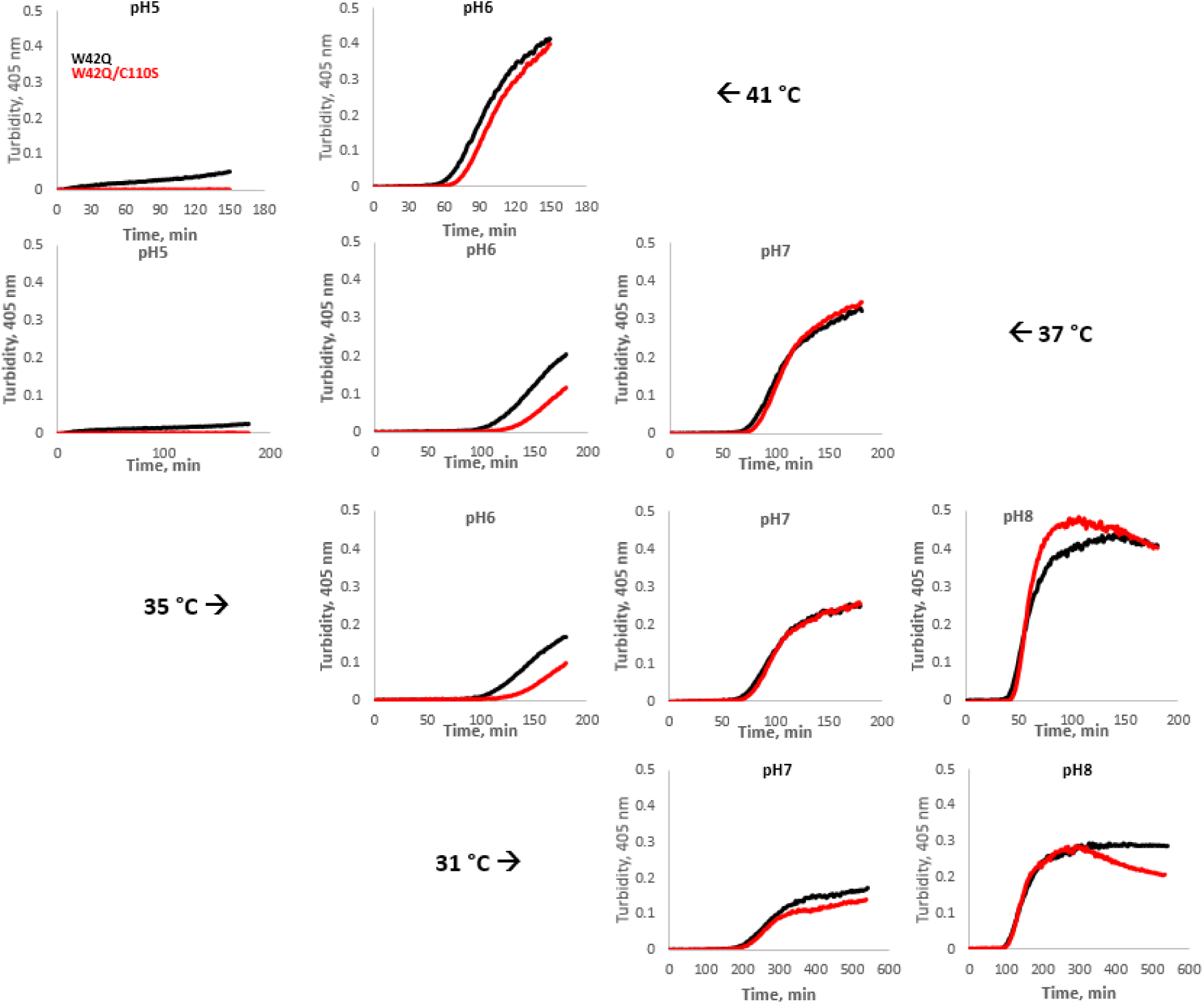
Temperature and pH dependence of WT(ox)-induced aggregation of W42Q (*black*) and W42Q/C110S (*red*) Representative turbidity traces are shown at temperatures of 31, 35, 37, and 41 °C and pH 5, 6, 7, or 8. All samples were buffered with phosphate, except pH 5, which was buffered with acetate. Note that the x-axes scales differ due to the slower aggregation kinetics at lower temperatures.

**Figure SI 5:**
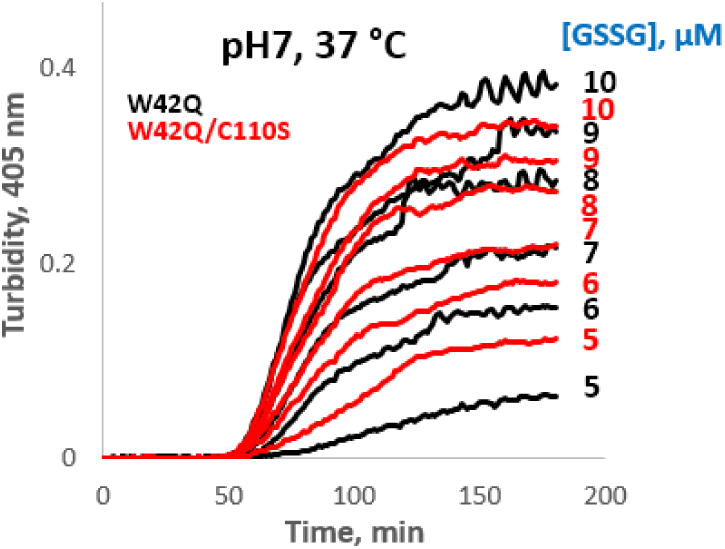
Effect from presence of Cys110 in the W42Q background depends on concentration of oxidizing agent. When the reduced W42Q (*black*) and W42Q/C110S (*red*) were incubated with various concentrations of oxidized glutathione, as indicated, no difference in aggregation lag time was observed. Presence of Cys110 appeared to be mildly protective, reducing turbidity amplitude, at the lowest concentrations of oxidizing agent, suggesting that formation of 108-SS-110 may serve as a decoy oxidation site under these conditions. By contrast, the W42Q/C110S mutant showed modestly less turbidity at the highest [GSSG], reversing this effect. A possible interpretation is that formation of the 108-SS-110 bond in the W42Q mutant leads to a modest destabilization of its N-terminal domain, thus exposing the Cys residues there to oxidation that traps aggregation-prone intermediates, such as the 32-SS-41 bond.

**Figure SI 6:**
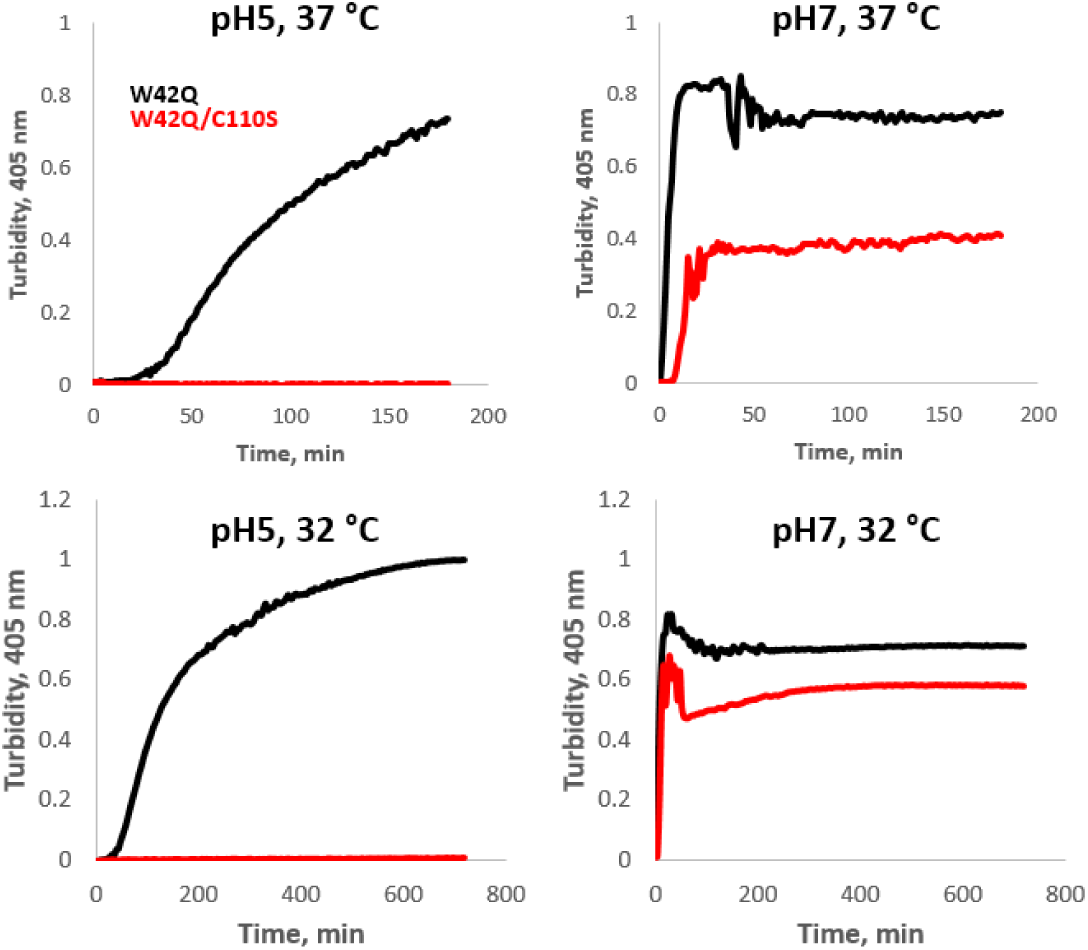
Cys110 is crucial for oxidative aggregation of the W42Q mutant at low pH. When reduced W42Q (*black*) and W42Q/C110S (*red*) were incubated directly with 1:3 copper(II):phenanthroline, at the concentration of 20 μM each of protein and copper(II), at pH5, only the W42Q variant aggregated. Note that this treatment produces 108-SS-110 bonds in the WT protein. However, at pH7 both variants aggregated rapidly and to a substantial degree, suggesting that other Cys residues were able to substitute for Cys110 at that pH. Kinetics of aggregation were temperature-dependent (note x-axes scale).

**Figure SI 7:**
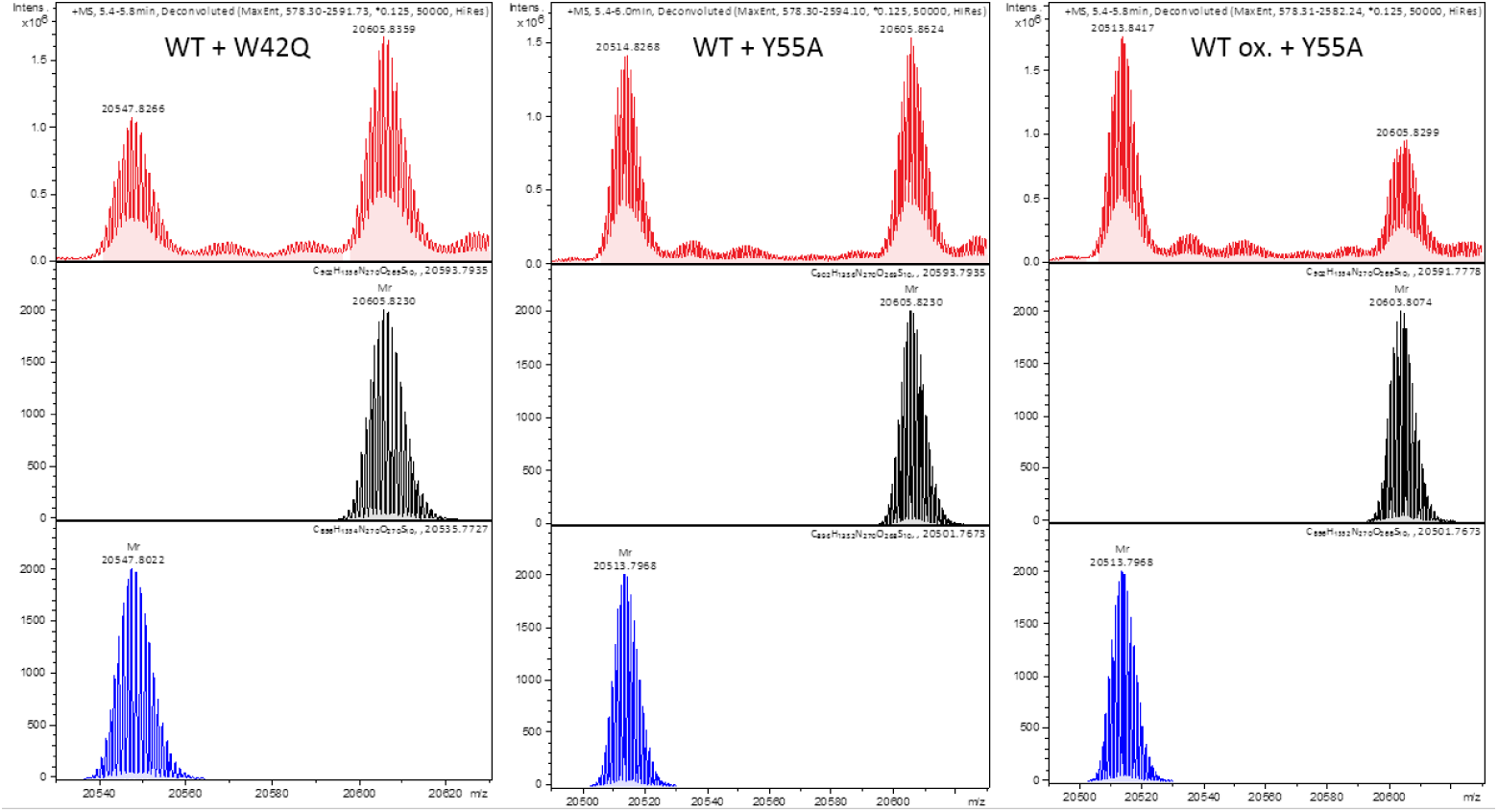
Mass spectrometric evidence for dynamic disulfide exchange in solution. Whole-protein isotopically resolved mass spectra (red) along with the corresponding predicted isotope distributions (black, blue) from control protein mixtures. In the mixture of reduced WT and reduced W42Q (left), both proteins retain their predicted isotope distributions at the end of the incubation, indicating that internal disulfides do not spontaneously form in any significant amounts. The same is true for a mixture of reduced WT and reduced Y55A, a soluble variant (center). However, when oxidized WT is mixed with reduced Y55A under the same conditions, the WT isotope distribution shifts to the right and the Y55A to the left, indicating some partitioning of the disulfide bonds between the two soluble variants.

**Figure SI 8:**
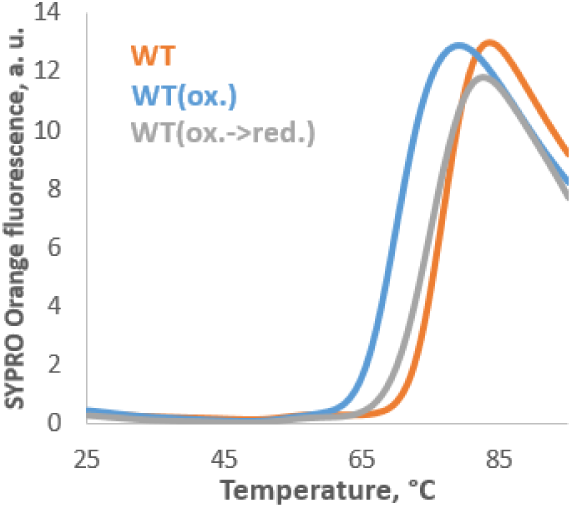
Differential scanning fluorometry of WT (*orange*), WT(ox.) (*blue*), and WT(ox.->red.) (*gray*), carried out at pH5 to inhibit any disulfide exchange, indicated virtually complete reversibility of the conformational destabilization introduced by the 108-SS-110 disulfide upon mild reduction (1 mM dithiothreitol for 1 h at 37 °C).

**Figure SI 9:**
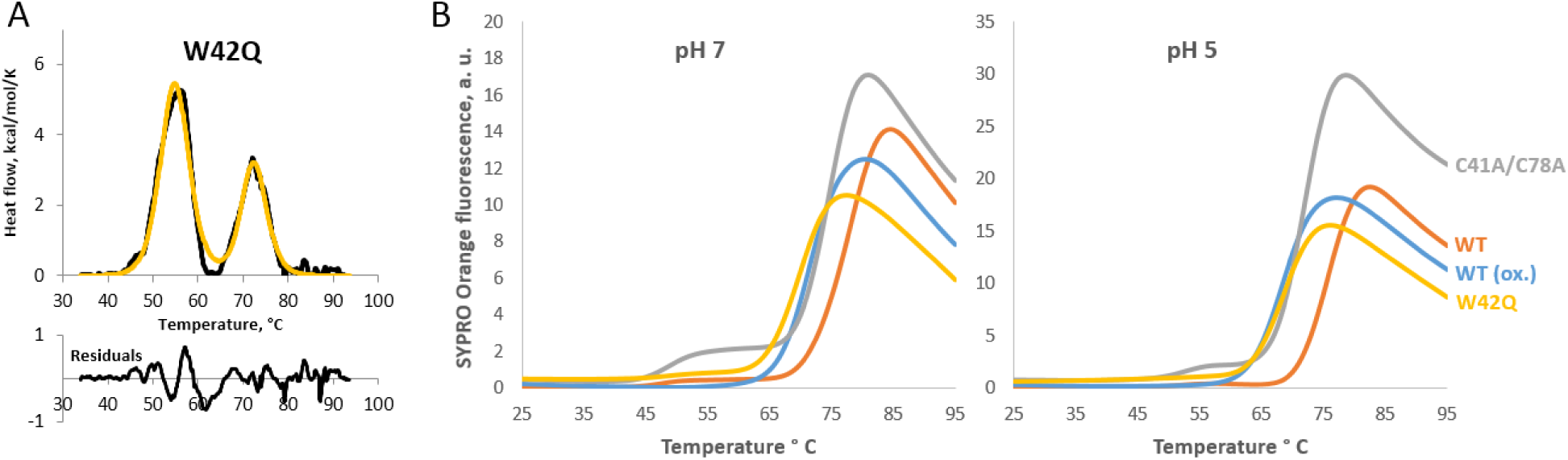
Differential scanning fluorometry reports on the foldedness of only the C-terminal domain. (*A*) Differential scanning calorimetry trace (*black*) and two-domain fitting (*yellow*) for the W42Q variant, from (Serebryany et al., 2016b). (*B*) Differential scanning flourometry of WT (*blue*), WT(ox.) (orange), W42Q (yellow), and C41A/C78A (gray) at pH 7 in 10 mM phosphate buffer and pH 5 in 20 mM acetate buffer. The lack of a detectable melting transition for the destabilized N-terminal domain of W42Q indicates that the SYPRO Orange reporter dye does not bind detectably to the molten globule state of the N-terminal domain.

**Figure SI 10:**
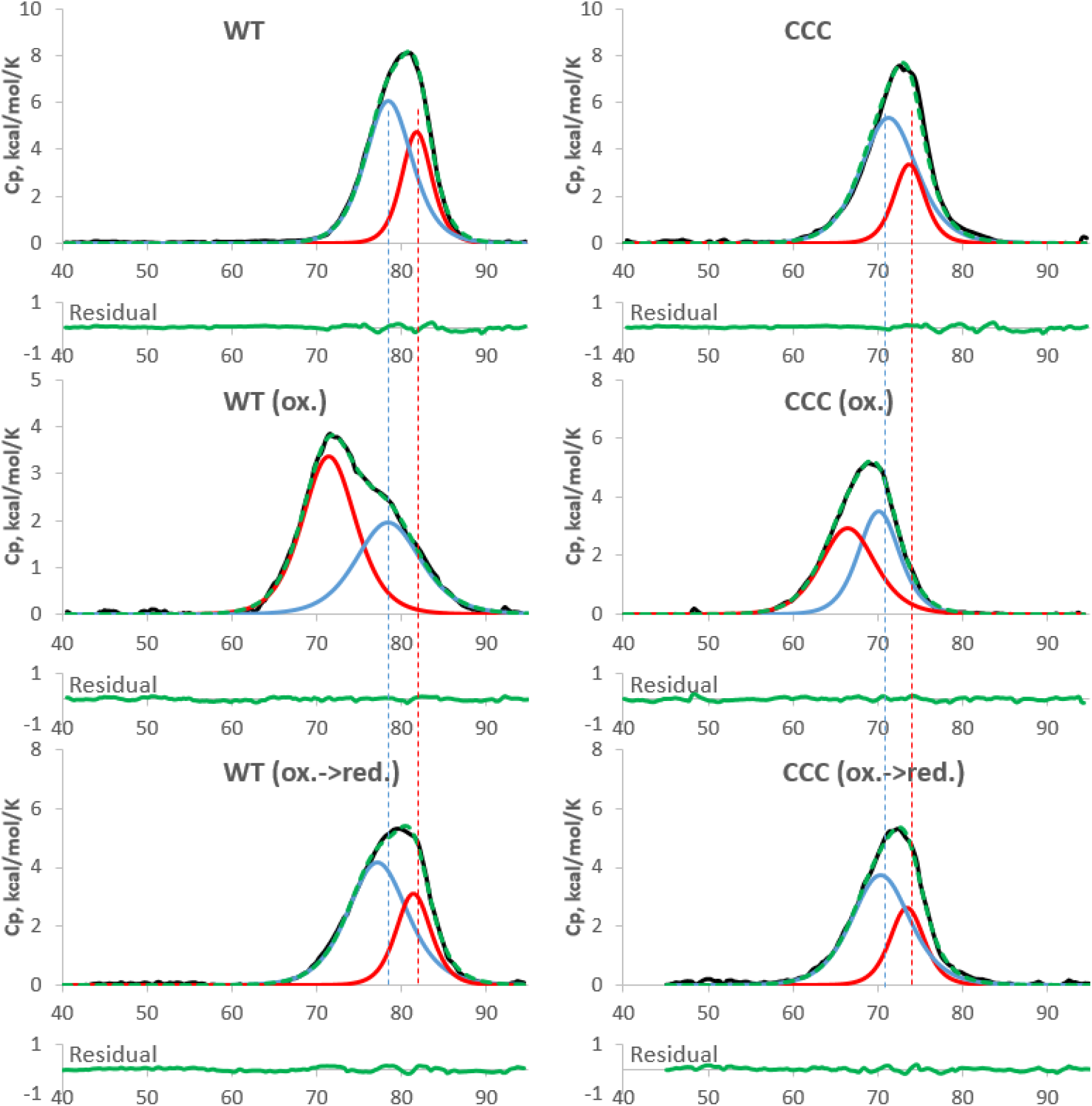
Differential scanning calorimetry thermograms of representative melts of the WT and C18T/C41A/C78A (“CCC”) HγD. Shown are: baseline-corrected thermograms (*black*); fitted melting transitions of the N-terminal (*blue*) and C-terminal (*red*) domains; sum of the fits (*dashed green*); and the fit residuals (*solid green*).

## Acknowledgments

This research was supported by the National Institutes of Health grant #GM111955 to E. I. S.

The authors thank Prof. Jonathan A. King for mentorship and discussion at the early stages of this research. We thank Renee Robinson for technical assistance with mass spectrometry.

